# Localized and Distributed Representations of Person Knowledge for Faces

**DOI:** 10.1101/2022.03.27.485948

**Authors:** Zarrar Shehzad, JohnMark Taylor, Gregory McCarthy

**Affiliations:** Department of Psychology, Yale University, New Haven, CT 06511; Department of Psychology, Harvard University, Cambridge, MA 02139

**Keywords:** anterior temporal lobe, fusiform face area, semantic information, classification, person-knowledge, face processing

## Abstract

Modular neural models of face processing posit that face-associated person-knowledge is localized in, or accessed through, the ventral anterior temporal lobe (vATL). However, some studies have suggested that person-knowledge is more widely distributed within a larger face-processing network that includes the mid-fusiform gyrus (‘fusiform face area’). Here, we conducted an fMRI study to distinguish whether person-knowledge is localized or distributed by comparing brain responses evoked by synthetic faces, for which participants had learned person-knowledge (biographical facts) and faces for which the subjects had learned only physical facts. After extensive training, participants were cued to recall a particular biographical or physical fact about the upcoming face. In an alternate passive-viewing task, participants were shown the same faces but were not cued to recall biographical or physical facts. Classification analyses (MVPA) were performed on several a *priori* chosen face-selective regions (ROIs) in the ventral temporal cortex. Within each ROI tested in isolation, MVPA discriminated faces associated with person-knowledge from faces only associated with physical facts. This result is consistent with a distributed model for person-knowledge. However, when tested in a single model to separate shared and unique information, unique information was limited to the left mid-fusiform and vATL posterior (vATL-p) ROIs. Moreover, the feature weights from these two areas showed that only left vATL-p was specialized for processing biographical facts. This latter result was obtained only when these biographical facts were explicitly retrieved in the cueing task. Thus, our results indicate that the left vATL-p represents explicit recall of face-associated person-knowledge.

**New & Noteworthy:** Whether person knowledge for faces is localized in a domain-specific region, vATL, or distributed in many domain-general brain regions, including the mid-fusiform gyrus, is hotly contested. We resolve this debate by using multivariate analyses to partial fMRI signal from different brain regions into unique and shared variance. Our findings show that unique information for person knowledge is found in both the vATL and mid-fusiform but only the vATL represents explicit recall of face-associated person knowledge.

---

Upon seeing a familiar face, we can easily access a wealth of associated biographical knowledge for that person. In many neural models of face processing, such biographical, or *person-knowledge*, is represented in the ventral anterior temporal lobe (vATL). The vATL has been suggested to link visual representations of an individual’s face with person-specific semantic knowledge (Collins and Olson 2014). The vATL is well positioned to support the association of visual and semantic knowledge, in that it has connections to ventral posterior occipito-temporal regions critical for face processing, and to medial and lateral temporal lobe regions that process semantics and memory (Moeller et al. 2008; Pyles et al. 2013; O’Neil et al. 2014). Indeed, the communication of information between the vATL and other brain areas might be as important in recognizing familiar faces as activity localized solely in the vATL.

Neuropsychological and neuroimaging evidence has provided support for a modular representation of face-associated biographical information within the vATL. Lesions to the ATL can cause an associative prosopagnosia, whereby individuals can perceptually discriminate between faces but have difficulty learning new information, or retrieving previously learned information, about a face (Damasio et al. 1990; Olson et al. 2007; Fox et al. 2008). However, lesions to the ATL that cause associative prosopagnosia can be extensive and often damage both its dorsal and ventral surface. Additional anatomical information regarding face processing in the ATL has come from electrical recordings in epileptic patients, where a face-specific event-related potential (ERP) recorded from subdural electrodes in the vATL was shown to be preferentially sensitive to name priming of faces (Nobre et al. 1994; Allison et al. 1999; McCarthy et al. 1999; Puce et al. 1999). These results are consistent with a meta-analysis of functional MRI (fMRI) studies showing that activity in the vATL was greater when viewing familiar/famous faces than novel faces (Von Der Heide et al. 2013). Moreover, responses to faces in the vATL are insensitive to many low-level perceptual manipulations that leave face identity intact such as contrast reversal (Nasr and Tootell 2012) and viewpoint changes (Anzellotti et al. 2014; Anzellotti and Caramazza 2016; Yang et al. 2016). Instead, responses in the vATL to faces are sensitive to prior knowledge such as pre-experimental familiarity (Eger et al. 2005; Rotshtein et al. 2005) or having learned a person's name and occupation (Tsukiura et al. 2010; Ross and Olson 2012). Taken together, these findings suggest that the vATL represents the identity associated with a person's face.

The vATL is part of an extended ‘face network’ (Haxby et al. 2000) that also includes two face sensitive regions in ventral occipito-temporal cortex (VOTC): the mid-fusiform gyrus (sometimes referred to as the fusiform face area, or FFA), and a region of cortex within and nearby the inferior occipital gyrus (IOG) (sometimes referred to as the occipital face area, or OFA). Some studies indicate that person-knowledge also depends upon the integrity of the face-sensitive mid-fusiform gyrus. For example, subdural electrical stimulation of the mid-fusiform gyrus in epileptic patients produces an inability to name famous and familiar faces (Allison et al. 1994; 1999). Similarly, lesions of the mid-fusiform result in difficulty learning the name associated with a face (Valdés-Sosa et al. 2011; Yang et al. 2016). A case study of an individual with an acquired prosopagnosia following damage to both his right FFA and OFA found that his right vATL showed repetition suppression to different images of the same face identity comparable to healthy controls (Yang et al. 2016). This suggests conserved functioning of the right vATL without any contributions from the right FFA and OFA. However, this individual still exhibited behavioral impairments in recognizing famous or familiar faces, which implies that activity from the mid-fusiform might be needed to access person-knowledge. Some functional magnetic resonance imaging (fMRI) studies have also reported activity in the mid-fusiform in addition to the vATL for faces learned in the laboratory and made familiar (Lehmann et al. 2004; Verosky et al. 2013). Finally, studies have also reported that patterns of brain activity in the mid-fusiform gyrus (in addition to the vATL) discriminate between face identities (Nestor et al. 2011; Verosky et al. 2013; Axelrod and Yovel 2015). These findings suggest that the vATL is not specific for representing biographical knowledge, and information might be distributed between at least the mid-fusiform and vATL.

Since these two areas are part of the face network and structurally connected, the information in each region is not independent (Haxby et al. 2000; Pyles et al. 2013; Collins and Olson 2014), and so activity in the mid-fusiform evoked by familiar faces could result from information originating in the vATL. The mid-fusiform, in this case, would not have any unique information for person knowledge but instead that information would have been shared by the vATL, resulting in the appearance of distributed representations. In a recent study (Shehzad and McCarthy, 2018), we developed an approach for distinguishing between unique and shared information and demonstrated the importance of this distinction for determining whether visual stimulus categories have localized or distributed neural representations. The typical approach taken in MVPA studies is to examine brain regions separately, and our results showed discriminable category representations in many brain regions indicative of a distributed representations when using this approach. However, MVPA applied in this manner does not distinguish between information in a region that is unique to that region versus shared among other regions. When we instead combined regions into a single model to remove shared information among regions, localized representations were revealed with unique information for each visual category localized to previously identified category-selective regions. Uniqueness of information hinges on how well a region can predict the desired outcome (e.g., biographical vs physical information) relative to other regions. Here, for the first time, we apply that same logic to determine if the mid-fusiform and vATL contribute unique information for representing person knowledge of faces.

Participants were trained over several days to associate initially novel synthetic faces with biographical facts (name, location, occupation) prior to testing in an fMRI experiment. To control for perceptual face familiarity, another set of faces were viewed the same number of times, but participants were trained on observable physical facts (eye color, gender, race). During a subsequent fMRI session, participants completed two tasks in which they viewed faces from each fact set condition, biographical (‘Bio’) or physical (‘Phys’).

In the Questions task, participants were visually cued to the condition (Bio or Phys) prior to each face presentation, and, after viewing the face, were asked if they knew a specific fact associated with the face. This task thus emphasized processing and explicit recall of person knowledge. In the Passive-Viewing task, participants simply viewed each face one at a time and were asked to reflect on what they knew about the face without making any explicit response. This task sought to assess the automaticity of processing person knowledge without the requirement for explicit recall. To identify unique and shared information, we followed Shehzad and McCarthy (2018) and measured in each task the classification accuracy for discriminating patterns of activity between Bio and Phys trials in each region separately and in a combined model.

Although our a *priori* hypotheses were directed towards the vATL and mid-fusiform, we extended our analyses to other functionally-defined face-selective brain regions that were identified in the Atlas of Social Agent Perception (ASAP) (Engell and McCarthy 2013). The vATL is not a precise anatomical term, and represents the confluence of many gyri. Two ROIs were identified in the ASAP as falling within the vATL, and we distinguish them as anterior (vATL-a) and posterior (vATL-p). The latter is located in the anterior-most fusiform gyrus. Although we are focused on the inferior temporal lobe, we included an ROI from the inferior occipital gyrus (IOG) which is typically denoted as the occipital face area, or OFA.

## Materials and Methods

### Participants

Sixteen right-handed, healthy adults (7 females, age range 18-25 years, mean 19.8 years) participated in the study. All had normal or corrected-to-normal vision and no history of neurological or psychiatric illnesses. The protocol was approved by the Yale Human Investigation Committee, and informed consent was obtained for all participants. Three additional participants completed the behavioral portion of the study, but did not reach the minimum criteria for memory of biographical facts and so did not participate in the fMRI study.

### Stimuli

Seventy-two synthetic faces were generated using the FaceGen software (Singular Inversions, Toronto, ON, Canada). We first created 36 faces with unique identities by systematically varying four features: gender (male, female), race (black, white, Asian), age (young, middle-aged, old), and facial expression (happy, sad), and then randomly varying the other features provided by FaceGen (e.g., eye size and facial structures). A second set of 36 faces was generated using the same process, and were matched to the first set for gender, race, age, and facial expression.

For one face set, three unique biographical facts were associated with each face: name, location (home state), and occupation (Figure 1). Names were chosen from a list of the most common names in 2013, states were randomly selected from 50 possible states in USA, and occupations were taken from a list of the top professions in the past decade (see Supplementary Table 1 [https://doi.org/10.5281/zenodo.3700452] for facts associated with each face). Each face of the second face set was associated with three physical facts: gender, eye-color, and race. Across participants, we counter-balanced the face set that was associated with the biographical or physical facts. For each participant, 36 faces were associated with biographical facts and 36 faces were associated with physical facts.

**Figure 1:**
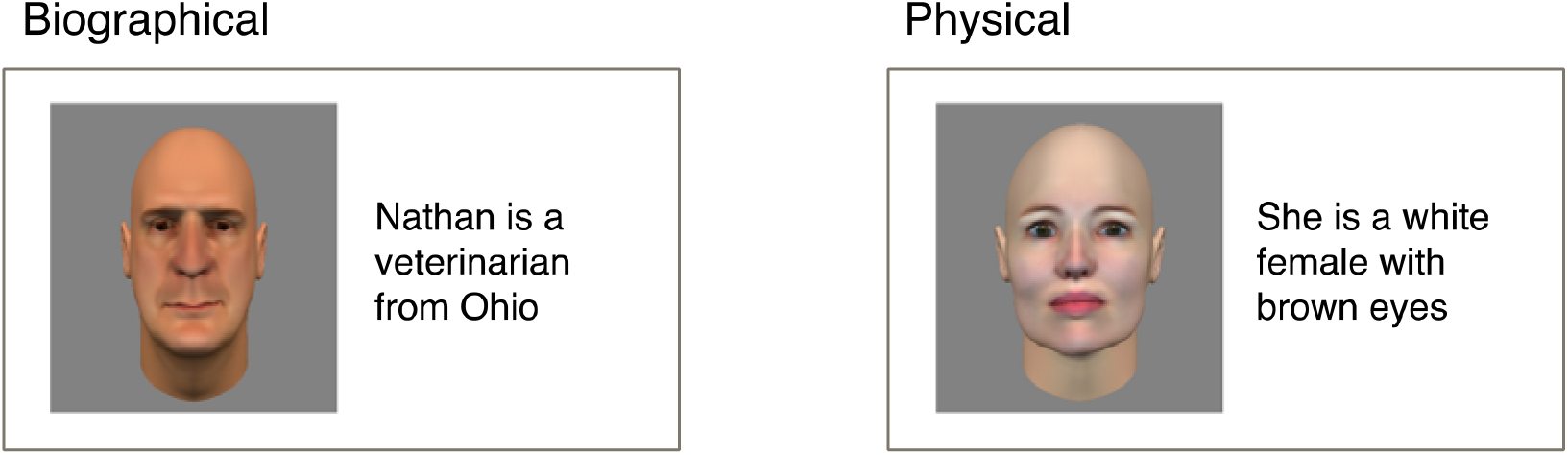
Sample training trial for faces associated with biographical facts (left) or physical facts (right) shown for 10s each.

### Procedure

The study comprised a minimum of five laboratory visits (Table 1). Participants were trained to associate each face with three facts (either biographical or physical) on three sessions occurring on consecutive days. On the first training day, but prior to training and when faces were all novel, an EEG data acquisition session was held. After training was completed, subjects viewed all faces in an fMRI session that generated the data presented herein. An EEG session was also held on another laboratory visit after the fMRI session. For three subjects, due to magnet scheduling issues, there was an average gap of 11 days between the training and fMRI session. For these subjects, an additional training session was given on the day preceding the fMRI scan.

**Table 1:** Listed are the experimental session number (first row) and activity during that session (second row). Each session was held on separate days typically only a day apart.

| Number | 1 | 2 | 3 | 4 | 5 |
| --- | --- | --- | --- | --- | --- |
| Activity | EEG, Training | Training | Training | fMRI | EEG |

The pre- and post-training EEG sessions were designed to investigate issues concerning the influence of perceptual familiarity and person-knowledge on particular event-related potentials (ERPs). This ERP study is now published (Taylor et al., 2016).

### Training Task

During each training session, face learning was separated into six blocks. In each block, participants were shown a face and the three associated facts for which they were required to remember for 10 s. Twelve faces, six with biographical facts and six with physical facts, were shown three times in random order within the block. Participants then saw each face from the current block with its category indicated (Bio or Phys) and were asked to verbally recall the facts associated with that face. The test was self-paced with a face remaining on the screen until the participant moved on to the next face. An experimenter noted the participant’s responses but did not provide immediate feedback. After testing, participants reviewed each face for 10 s with the correct facts listed alongside. When all six blocks were completed in this manner, participants were tested on all 72 faces and were asked to recall the facts associated with each face as before. Participants were required to correctly recall more than 40 of 108 biographical facts at the end of the first session in order to continue in the experiment.

### fMRI Tasks

Participants completed two tasks during the functional magnetic resonance imaging (fMRI) session. The primary *Questions* task actively engaged participants in the memory of a face by requiring them to retrieve facts associated with each face. A second, *Passive Viewing* task was also included because many of our and other’s prior face ERP and fMRI studies used passive viewing designs, and we were curious to see whether activation differences found in an active memory retrieval task were evident in the absence of explicit cues to recall biographical information.

In the *Questions* task, participants viewed each of the 72 faces seen during training (36 Bio and 36 Phys). Each trial began with a word cue indicating whether the upcoming face belonged to the Bio or Phys condition. This condition cue was displayed for 1 s. A face from that fact set immediately followed the cue and was displayed for 4 s. At face offset, a probe question appeared for 4 s that specified one fact appropriate for the fact set. Participants pressed one of two buttons to indicate whether they knew the fact or not. A fixation-cross then appeared for an inter-trial interval (ITI) of 8, 10, 12 s. Faces were presented in random order over four runs. Each run consisted of 18 trials lasting 263 s.

Participants also completed a secondary *Passive Viewing* task. Here participants viewed the same 72 faces as seen in the Questions task and were instructed simply to reflect about what they knew about the face without making an overt response. Faces appeared for 4 s with an inter-trial interval (ITI) of 13, 15, or 17 s during which a fixation-cross was displayed. Faces were presented in random order over four runs. Each run consisted of 18 trials lasting 263 s. The runs alternated between the Passive Viewing task, which was always first, and the Questions task. All stimuli were presented on the center of a screen (10° × 10°) located behind the participant in the scanner and viewed with a mirror mounted in the head coil.

#### fMRI Memory Test

After completing the fMRI scan, participants were given a surprise memory test for those faces for which they learned biographical information. Participants were shown each face and asked to remember all three facts associated with each face, just as they did at the end of each training session. Responses were given verbally with no time limit.

### Behavioral Analysis

All behavioral analyses on memory performance and response time were conducted in the R statistical package (version 3.1.2). We conducted a two-way repeated measures analysis of variance (ANOVA) on the memory performance for each biographical fact recalled at the end of every session (three training and one MRI). For the response time, we conducted a one-way

ANOVA examining the main effect of condition (Bio or Phys) on response time. Additional one-way ANOVAs were conducted for each condition to examine the main effect of physical facts on response time and the main effects of biographical facts on response time.

### Image Acquisition and Preprocessing

Data were acquired using a 3T Siemens TIM Trio scanner with a 32-channel head coil. Functional images were acquired using a multiband echo-planar pulse sequence (TR = 1000 ms, TE = 32 ms, flip angle = 62°, acceleration factor = 3, FOV = 210 × 210 mm, matrix = 84 × 84, slice thickness = 2.5 mm, 52 slices, voxel size = 2.5 mm^3^). A high-resolution structural image was acquired for registration using a 3D MP-RAGE sequence (TR = 2530 ms, TE = 2.77 ms, flip angle = 7°, FOV = 256 × 256 mm, matrix = 256 × 256, slice thickness = 1 mm, 176 slices).

Image preprocessing was performed using custom scripts (available on http://github.com/HumanNeuroscienceLab/facemem) that incorporate functions from AFNI (v 2014-10-23 OpenMP, http://afni.nimh.nih.gov/afni), FSL (v 5.0.7, http://www.fmrib.ox.ac.uk/fsl), and Freesurfer (v 5.3.0, http://surfer.nmr.mgh.harvard.edu). The pipeline for functional images proceeded as follows: (1) the first 6 volumes (6 s) were discarded to allow for MR equilibration, (2) motion correction was performed on the functional images using AFNI’s 3dvolreg, (3) skull stripping of the mean functional image was performed using FSL’s BET (Smith 2002), (4) spatial smoothing with a 5-mm FWHM Gaussian mask was performed using FSL’s SUSAN (Smith and Brady 1997), (5) high-pass filtering with a 0.01 Hz cut-off to remove low-frequency drift was conducted using FSL, as was mean-based global intensity normalization. Structural images were skull-stripped with a hybrid watershed/surface deformation procedure using Freesurfer (Ségonne et al. 2004). The functional images were registered to the high-resolution structural images with boundary-based registration using FSL’s Flirt. The structural images were in turn nonlinearly registered to the Montreal Neurological Institute’s MNI152 template (2 mm isotropic) using FSL’s FNIRT (Andersson et al. 2007) and this transform was then applied to the functional images in anatomical space.

### Regions of Interest

We generated regions of interest (ROIs) to examine the temporal response to each condition and classification between conditions in face-selective regions. We identified regions of interest in the inferior occipital gyrus (IOG) (typically called the occipital face area, or OFA), the mid fusiform gyrus (mFus) (typically called the fusiform face area, or the FFA), and ventral anterior temporal lobe (vATL) based on the Atlas of Social Agent Perception (Engell and McCarthy 2013). We used the probability maps of face-selective responses from two localizers (face vs. house and face vs. scene) found in the atlas. The probability maps represented the percentage of participants who showed activation (p < 0.05, one-tailed) for the face-selective contrast. To identify activation peaks, we first averaged the two probability maps (face > house and face > scene), then smoothed the map by 2mm using AFNI’s 3dBlurInMask, and finally used AFNI’s 3dExtrema to find the local maxima (minimum distance between peaks = 12 mm; probability threshold = 0.1). We selected those local maxima found in the fusiform gyrus and lateral occipital cortex based on Freesurfer’s cortical labeling of the MNI152 brain. Eight peak locations were found, including the left inferior occipital gyrus (IOG; X = −44, Y = −82, Z = −10; BA 18), left mid-fusiform (mFus; X = −40, Y = −50, Z = −18; BA 37), left vATL posterior subregion (vATL-p; X = −42, Y = −28, Z = −20; BA 20), left vATL anterior subregion (vATL-a; X = −34, Y = −6, Z = −34; BA 20), right IOG (X = 42, Y = −78, Z = −10; BA 18), right mFus (X = 42, Y = −50, Z = −20; BA 37), right vATL-p (X = 44, Y = −28, Z = −22; BA 20), and right vATL-a (X = −34, Y = −6, Z = −34; BA 20).

For each peak, we created an ROI with a sphere of 4mm radius in MNI152 space (2 mm isotropic) using AFNI’s 3dUndump, which were then transformed into each participant’s functional space (2.5 mm isotropic). The total size of each resulting ROI was then 33 voxels or 264mm^3^. The average probability for face activity (average of face > house and face > scene) in each ROI was 0.37 for the right IOG, 0.45 for the right mFus, 0.12 for the right vATL-p, and 0.18 for the right vATL-a, 0.40 for the left IOG, 0.10 for the left mFus, 0.01 for the left vATL-p, and 0.09 for the left vATL-a.

### Univariate whole-brain analysis

Our issues of interest in this paper are addressed by the multivariate analyses described below. Nevertheless, for continuity with the extant literature and to provide context to the multivariate results, we conducted whole brain (Bio > Phys) contrasts using the standard general linear model approach and also examined the average temporal response in our face-selective ROIs.

#### Voxel-wise analyses

Whole-brain voxel-wise regression analyses were performed at the subject-level using AFNI’s 3dDeconvolve and 3dREMLfit functions, which provide temporal pre-whitening via an autoregressive model. The linear model consisted of two explanatory variables, one for trials when participants viewed faces associated with biographical facts (Bio condition) and another for trials when participants viewed faces associated with physical facts (Phys condition). All variables were modeled as boxcar functions, where a value of 1 was assigned to time-points when the face was on the screen (4-s per trial) and a value of 0 to time-points when the face was not on the screen, and then convolved with a double-gamma function (AFNI’s SPMG1). Regressors of no interest included baseline effects of each run and 6 head movement parameters. Contrasts were defined to identity brain regions that showed increased activity when viewing faces associated with biographical versus physical facts (Bio > Phys) or the inverse (Phys > Bio). Following individual analyses, whole-brain group analysis was performed using AFNI’s 3dMEMA (Chen et al. 2012), a program that incorporates individual beta precision estimates into group effects using a mixed-effects meta-analytic approach. Clusters were defined as contiguous sets of voxels with Z > 1.96 and then thresholded using Gaussian random field theory (cluster probability p < 0.05) to correct for multiple comparisons (Worsley et al. 1996).

#### Average evoked time-series

We took the mean time-series within each ROI from the probabilistic atlas and overlap in activity. The data for the time-series were prepared by concatenating the four runs in each task and regressing out the run effects and six motion parameters. Epochs time-locked to the onset of a face presentation were then extracted from the continuous time-series of a ROI, and the mean signal for the three time-points preceding onset was subtracted from signal at every time-point in the epoch. To smooth the data, the 36 epochs for each trial type (faces associated with biological facts or physical facts) were fitted with the general additive model using smoothed splines via the R package *mgcv* (Wood 2011). This resulted in smoothed average epochs that were 24 secs long (TR = 1 sec) and consisted of 5 secs preceding and 19 secs following the onset of the face on the screen. Smoothed average epochs were collected at each ROI. To get estimates of the time to peak or peak latency, we found the time-point with the maximum value in each smoothed average epoch.

### Classification or Multivoxel Pattern Analysis (MVPA)

The univariate analyses considers each voxel separately and cannot assess the nature of the neural information for person knowledge. Is the information for person knowledge in the mid-fusiform and vATL shared between regions or unique to each region? To address this question, we used classification analyses to determine if biographical information is distributed across face-selective regions or localized in particular regions.

In each of the ROIs from the probabilistic atlas, we tested the amount of information for discriminating biographical knowledge using MVPA (Bio > Phys). To perform the pattern analysis, we obtained parameter estimates (or betas) for each trial at the subject-level using AFNI’s 3dDeconvolve and 3dREMLfit functions. We applied a linear model with 72 task regressors (one for each trial; 36 for the Bio condition and 36 for the Phys condition) and additional covariates of non-interest, including baseline effects for each run and 6 motion parameters. Each task regressor was modeled using a double-gamma function with a duration of 4s and produced a volume of voxelwise betas. We concatenated the volumes into a single beta series for each condition (biographical or physical) and then extracted the beta series at every voxel in each ROI. We repeated this process for each participant and each task.

A two-way classification (biographical vs physical condition) based on a logistic regression was performed on the betas in each ROI using the elastic net, as implemented in the R packages caret (Kuhn 2008) and glmnet (Friedman et al. 2010). The elastic net is a regularized regression method that can account for collinearities in high-dimensional data and is a blend of the L1 (lasso) and L2 (ridge) penalties. Each voxel’s beta series was Z-scored before entry in the elastic net model to allow for the model to converge as well as account for differences in activity levels between voxels. Classification training and testing were performed using a leave-one-run-out cross-validation strategy. Group-level classification accuracies were obtained by averaging the accuracies from all participants for each task. Significance was assessed with a permutation test. For each subject, random permutations were applied to the trial indices 500 times and the classification accuracies were re-computed each time. The null distribution was simulated by calculating the group-level classification accuracies at each permutation. The group-level classification accuracy from the original data was then referred to this simulated distribution to obtain a p-value. Confusion matrices were also generated during classification to assess if the two-way classification discriminated both conditions successfully instead of just one condition.

We explored the amount of unique information for biographical knowledge among our ventral surface ROIs by comparing the classification accuracy for a “macro-ROI” consisting of the union of all ROIs, against a macro-ROI consisting of the union of all ROIs except for a target ROI. A high value indicates that the target ROI contributes additional (unique) information over and above all the other ROIs towards discriminating biographical and physical knowledge. Since there are considerable redundancies between the two hemispheres, we ran this analysis separately for the ROIs within each hemisphere.

Since the classification accuracy merely tells us there is a difference between the two conditions, we examined the feature weights to understand the direction of this difference (i.e., are activity patterns more related to the biographical or physical condition). Feature weights were extracted for each subject and task from our multivariate logistic regression that included all regions together in each hemisphere. In this model, the weights at each voxel reflect the contribution of that voxel after partialling out activity from all other voxels in the same hemisphere (i.e., the amount of unique information at that voxel). As a comparison, we also used feature weights from a mass-univariate logistic regression (i.e., with the regression run on one voxel at a time). For both univariate and multivariate analyses, we counted the number of weights that were negative or positive. For negative weights, an increase in activity at that voxel was associated with a higher likelihood of the biographical condition (Bio > Phys), whereas for positive weights, an increase in activity at that voxel was associated with a higher likelihood of the physical condition (Phys > Bio). In this way, we interpret the weights from our multivariate analysis in a manner similar to interpreting activity levels from a univariate analysis (Shehzad and McCarthy, 2018). A voxel with a positive weight associated with the physical condition could be used to predict the presence of person knowledge (biographical condition) simply because the trial activity is NOT from the physical condition. In an extreme case if all the voxels in a region have a positive weight, then such a region would have high classification accuracy for discriminating between the biographical and physical conditions but would contain information solely associated with perceptual knowledge (physical condition) rather than person knowledge (biographical condition). Consequently, stronger evidence for neural representations related to person knowledge would be found in a region having more voxels with negative weights associated with the biographical condition.

## Results

Here we report the behavioral performance, univariate analyses, and classification analyses that examine the multivariate representations for person knowledge along the ventral surface.

### Behavior

Table 2 presents descriptive statistics for memory performance of each biographical fact (location, name, occupation) recalled at the end of each session (three training and one MRI). We found a significant main effect of session (F[1,171] = 124.6, *p* < 0.05) but no significant effect for the type of biographical fact (F[2,186] = 0.8, *p* = 0.47). Memory performance increased across sessions and reached a plateau by the end of the third session (93% ± 14% of facts were remembered, mean ± standard deviation across participants).

**Table 2:** The percent of biographical facts recalled at the end of each session are shown below (mean ± standard deviation taken across subjects).

| Session | Location | Name | Occupation | Total |
| --- | --- | --- | --- | --- |
| *1 - Training* | $55 \pm 16$ | $55 \pm 16$ | $60 \pm 15$ | $57 \pm 15$ |
| *2 - Training* | $85 \pm 15$ | $84 \pm 17$ | $87 \pm 12$ | $85 \pm 14$ |
| *3 - Training* | $93 \pm 15$ | $92 \pm 15$ | $94 \pm 11$ | $93 \pm 14$ |
| *4 - MRI* | $93 \pm 14$ | $91 \pm 16$ | $95 \pm 11$ | $93 \pm 14$ |

In the MRI scan, response time (RT) was measured to a probe question for the biographical and physical condition. There were no significant effect of condition on response time (F[1,79] = 1.7, p = 0.2). Furthermore, within each condition, there was no significant effect for the particular fact probed (biographical condition: F[2,30] = 1.0, p = 0.37; physical condition: F[2,30] = 0.3, p = 0.76). We did observe on average slightly longer response times during the biographical condition (M = 1.43s ± 0.66s) than the physical condition (M = 1.35s ± 0.45s).

### Univariate (GLM) Task Analysis

#### Questions Task

The Bio > Phys contrast evoked a cluster of activation centered in the left vATL-a and peaking in the left vATL-p with few significant voxels in the left mFus (Fig 2). Results were predominately left-lateralized and included brain areas involved in attention and memory such as the retrosplenial cortex, parahippocampal gyrus, caudal anterior cingulate, and anterior insula (Red-Yellow in Fig 3). The reverse Phys > Bio contrast revealed less extensive differences with significant voxels within auditory, motor, and somatosensory cortices (Blue-Cyan in Fig 2). Coordinates of all local maxima are reported in Table 3. To drill down into the GLM results, we examined the temporal response in face-selective regions in the IOG, mFus, vATL-p, and vATL-a (Supp Fig 2). We found that activity during the biographical condition was significantly greater than the physical condition in the left vATL-p between 3.4s and 10s following trial onset (all *ps* < 0.05, paired t-test at each time-point; Supp Fig 2b). In contrast, none of the other ROIs had a significant difference between conditions at any time-point (all *ps* > 0.05). Taking the area under the curve (AUC) for each temporal response corroborated these findings wherein greater activity (AUC) was found in the Bio versus Phys condition for the left vATL-p (t[15] = 2.2, *p* < 0.05).

**Table 3:**
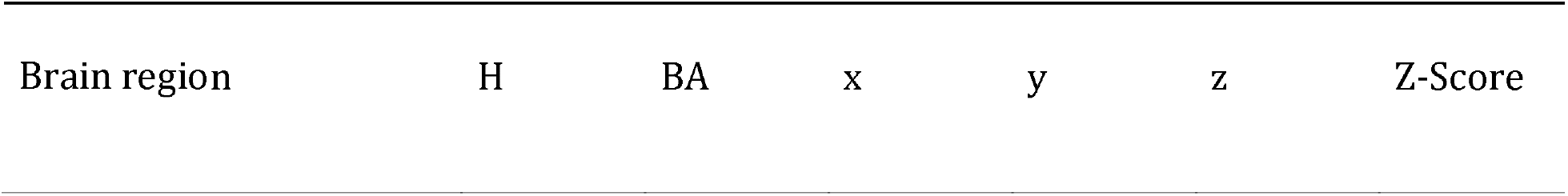
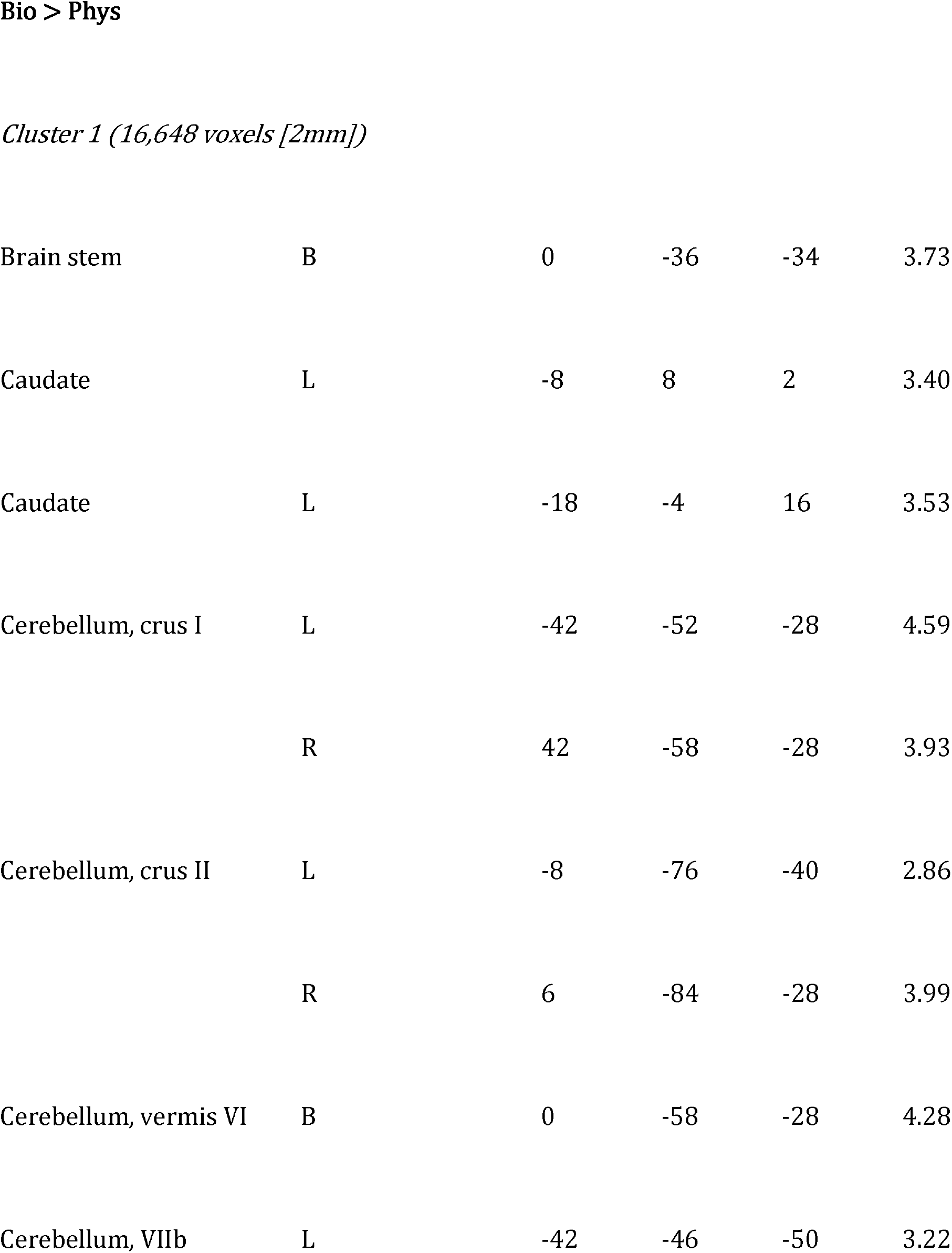
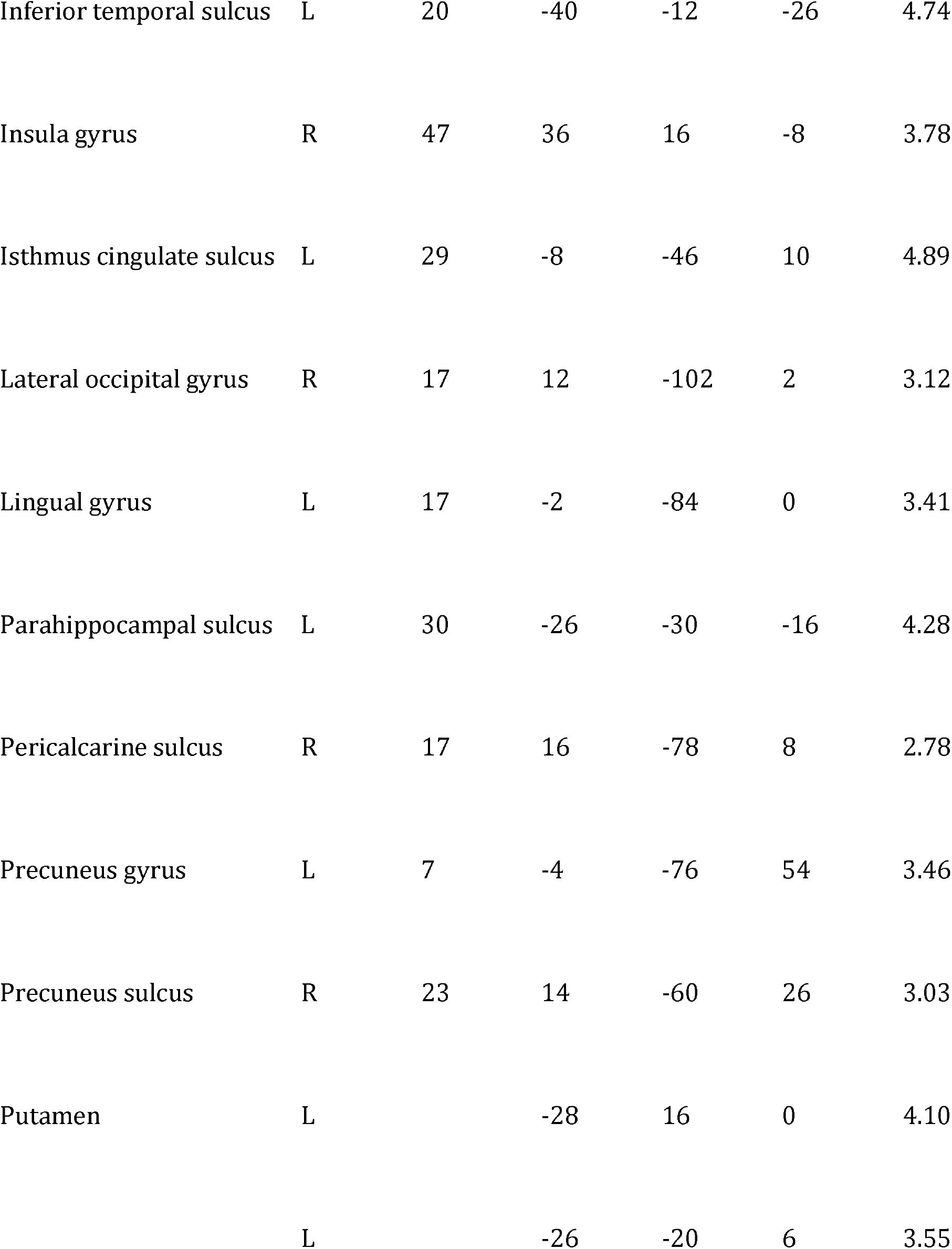
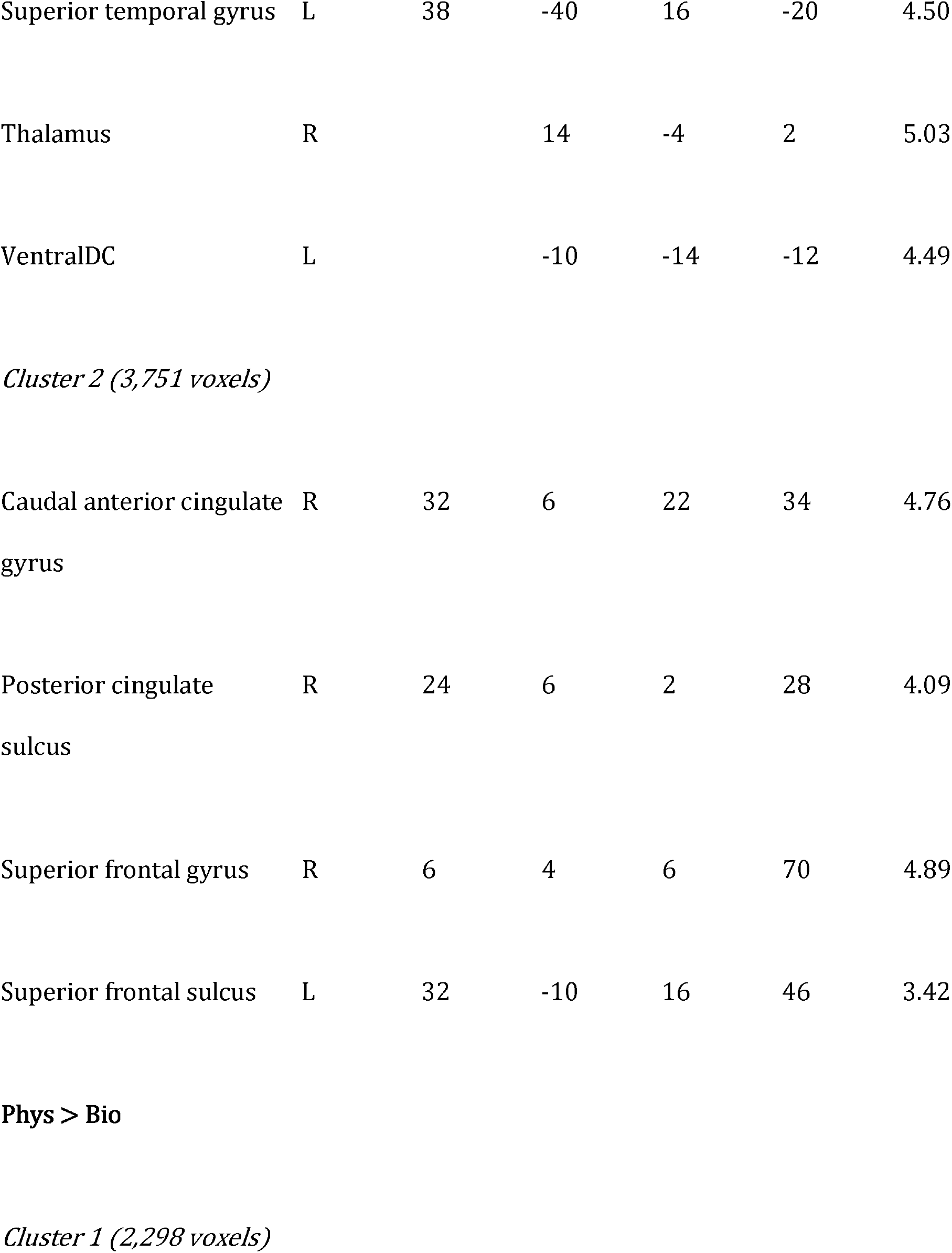
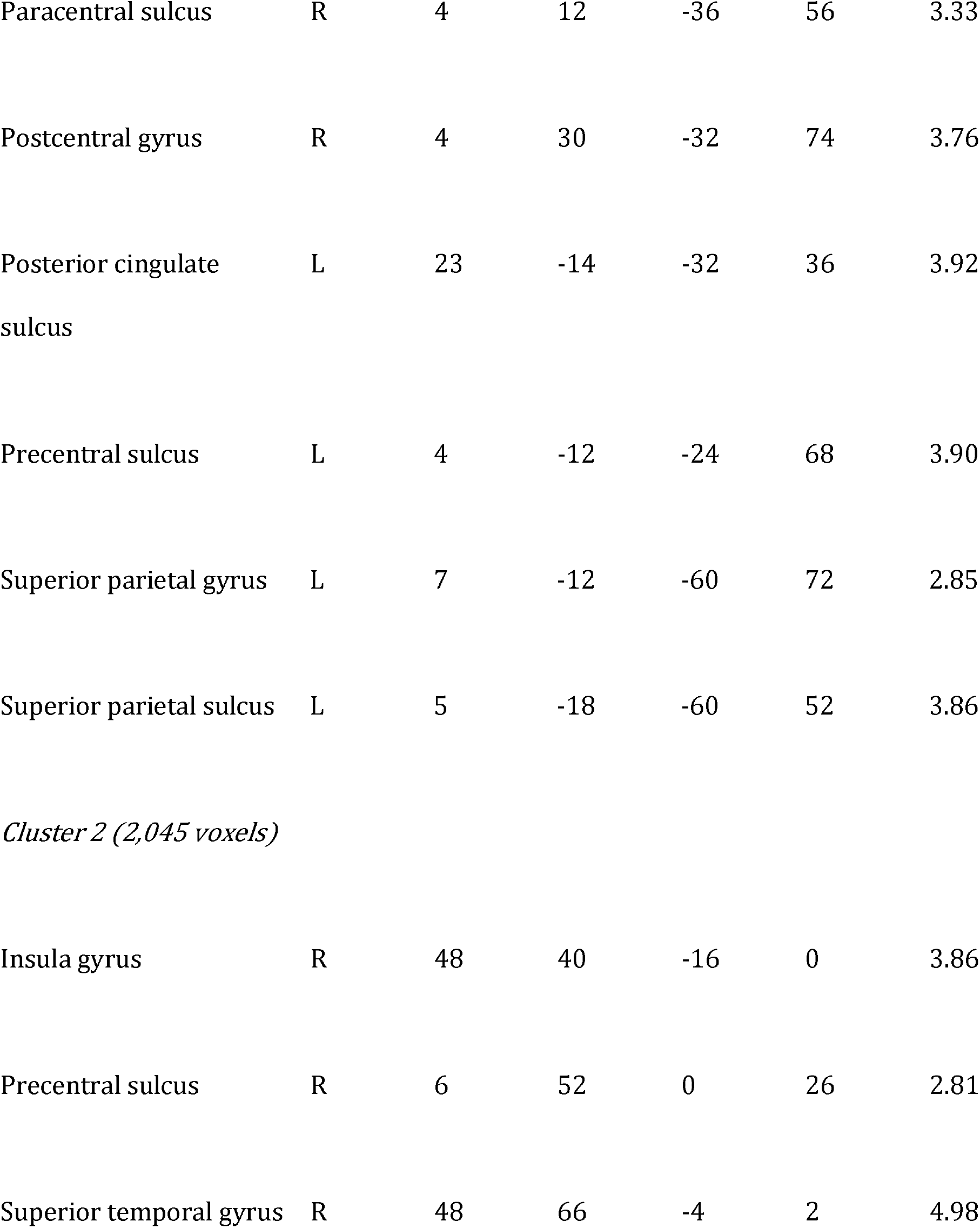
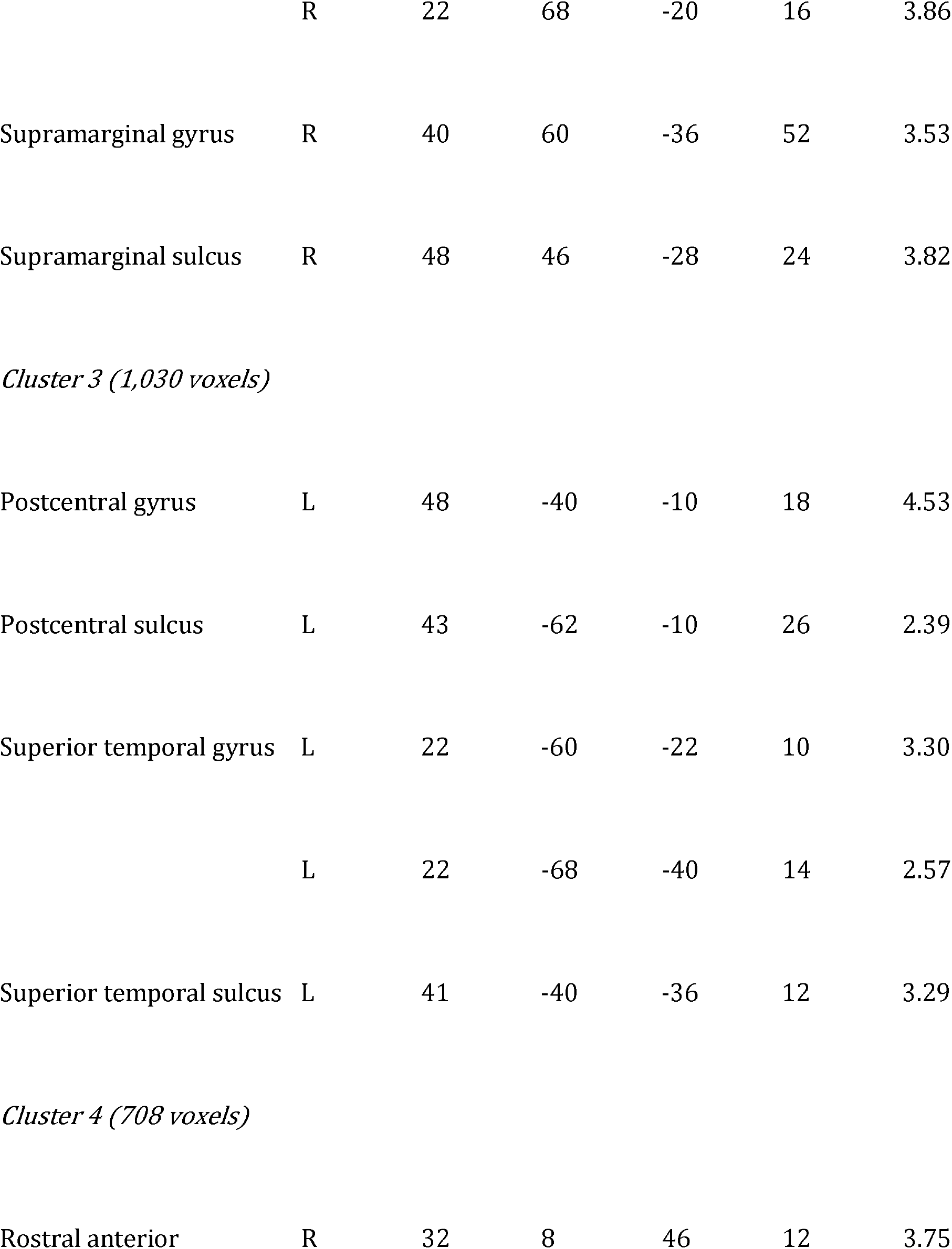
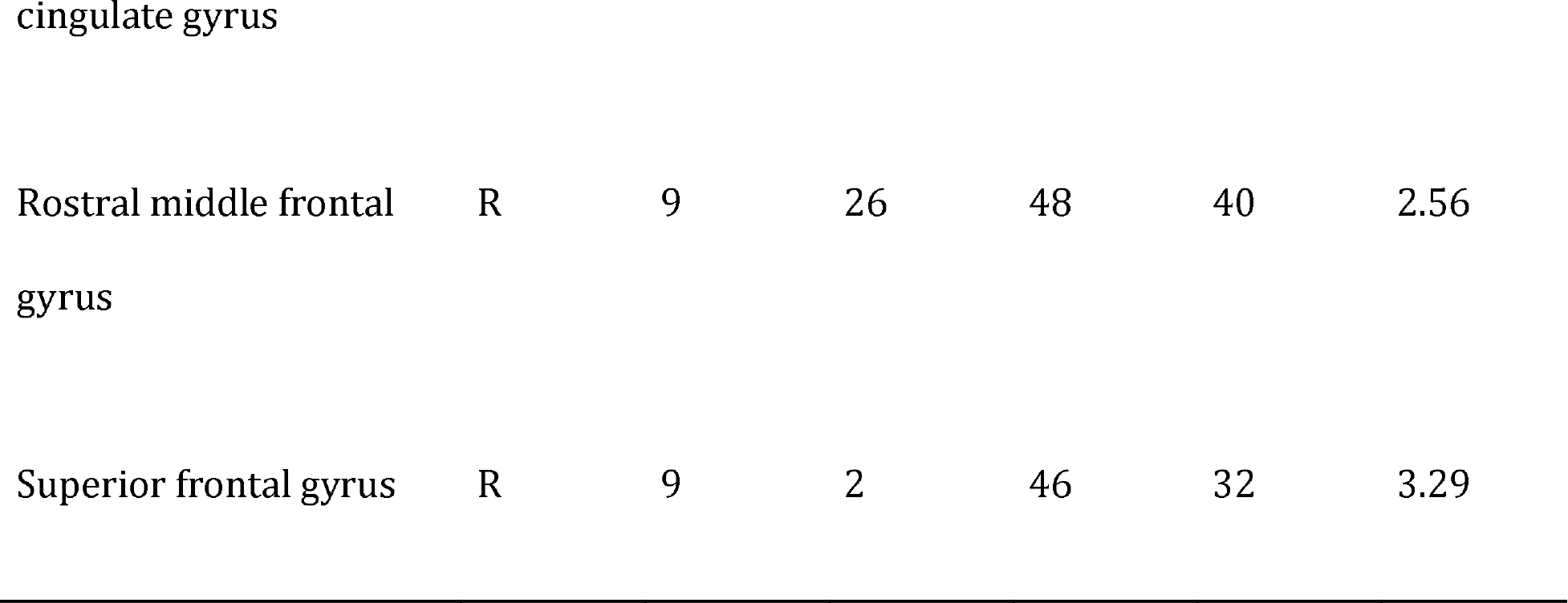
Listed are the peaks of significant brain activity for faces associated with (bio)graphical versus (phys)ical information. Peaks were found using AFNI's 3dExtrema by setting a voxel-level threshold of Z > 1.96 (cluster corrected, p < 0.05) and a minimum distance of 20 mm between peaks. Abbreviations: H, hemisphere; B, bilateral; R, right; L, left; BA = Brodmann Area.

**Figure 2:**
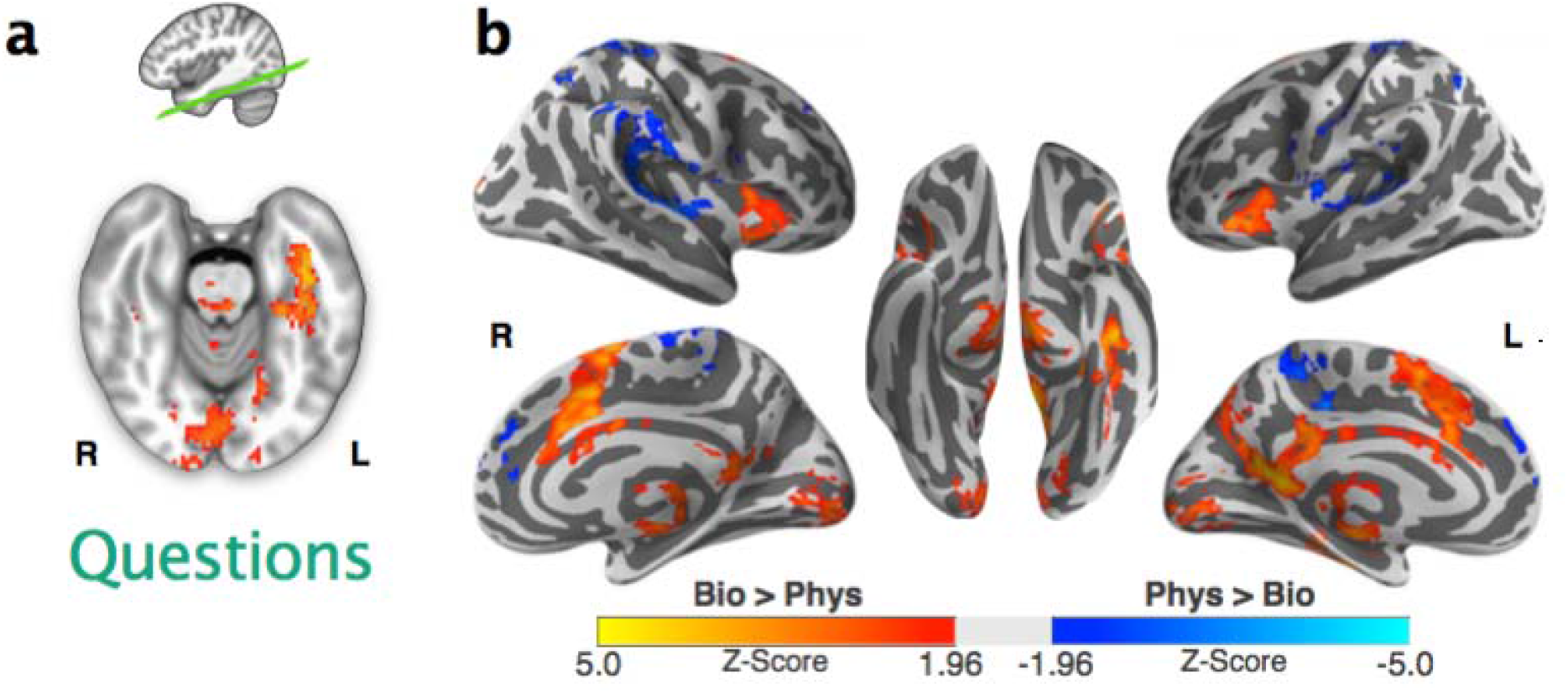
Difference in activity between the biographical and physical conditions for the Questions task (*p* < 0.05, cluster-corrected). Significantly greater activity for the biographical versus physical condition (red-yellow) and physical versus biographical condition (blue-cyan) is shown on (a) an oblique axial slice in MNI152 space (Z = 24 with 20 degrees rotation) and (b) on the fsaverage surface.

**Figure 3:**
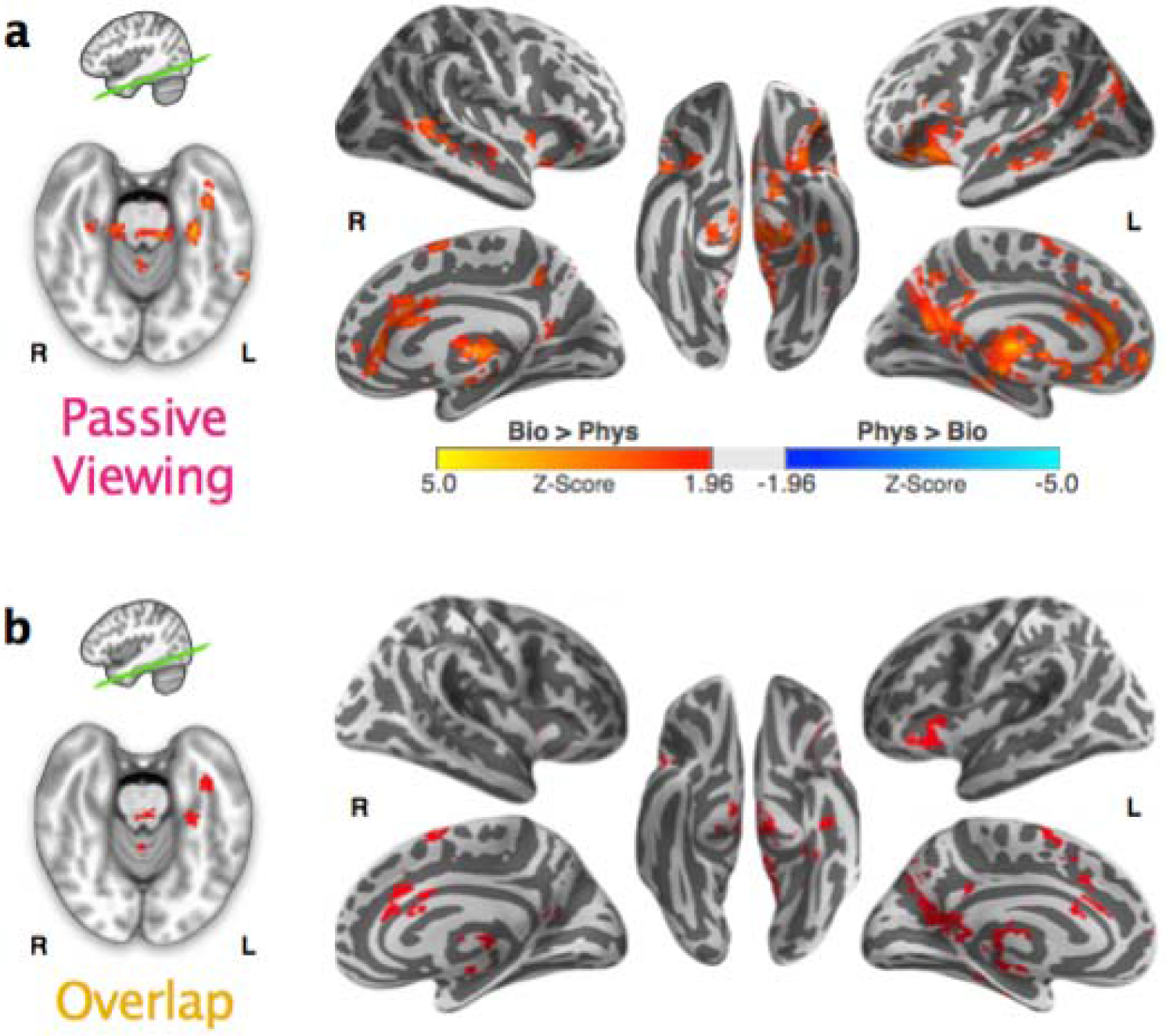
(a) Difference in activity between the biographical and physical conditions for the Passive Viewing task (*p* < 0.05, cluster-corrected). Significantly greater activity for the biographical versus physical condition (red-yellow) and physical versus biographical condition (blue-cyan; no voxels) is shown on an oblique axial slice in MNI152 space (Z = 24 with 20 degrees rotation) and on the fsaverage surface. (b) Overlap in activity for the Bio > Phys contrast between the Question and Passive Viewing task is shown in red.

#### Passive Viewing

In the left vATL (peaking in the vATL-p), we found significantly greater activity during the Bio condition than the Phys condition, which was similar to the Questions task (Fig 3a). However, unlike the Questions task, the Bio > Phys response during Passive Viewing was more focused on the left vATL and there were fewer significant effects extending posteriorly to the left mid-fusiform. When comparing the Passive Viewing and Questions task, the pattern of brain response for the unthresholded Bio > Phys contrast was similar between the two tasks (Spearman Rho = 0.39; dice coefficient = 0.36). Brain regions that showed significant activity (Bio > Phys) in both tasks tended to be left-lateralized and included cortical regions (retrosplenial cortex, parahippocampal gyrus, caudal anterior cingulate, and anterior insula) and subcortical regions (thalamus and caudate) (Fig 3b). Regions that had significant activity (Bio > Phys) in the Passive Viewing task but not Questions task included frontal ventromedial cortex as well as clusters along the superior and middle temporal gyri. There were no significant results for the Phys > Bio contrast. Coordinates of all local maxima are reported in Table 4. There was no significant difference in the temporal response between conditions at any time-point for any of the ROIs (all *ps* > 0.05, corrected; Supp Fig 3). In addition, no differences were found between conditions in the AUC (all *ps* > 0.2), and no differences between conditions were found in the peak latencies (all *ps* > 0.16). However, the temporal response for the Passive Viewing task showed a similar profile to the Questions task with the Passive Viewing task having less pronounced differences between the Bio versus Phys condition especially for the vATL-p and vATL-a (mean Spearman Rho = 0.69 ± 0.2).

**Table 4:**
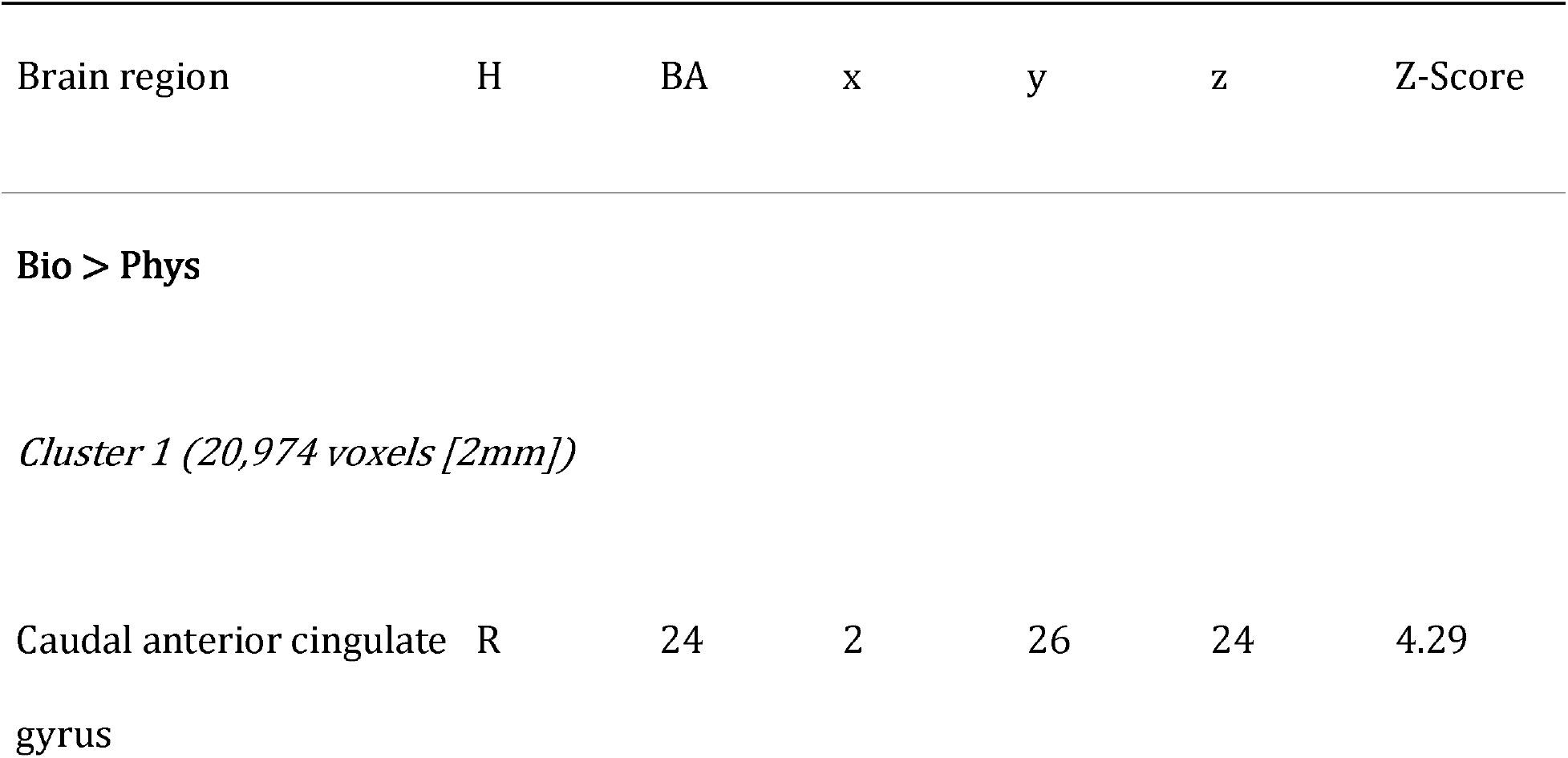
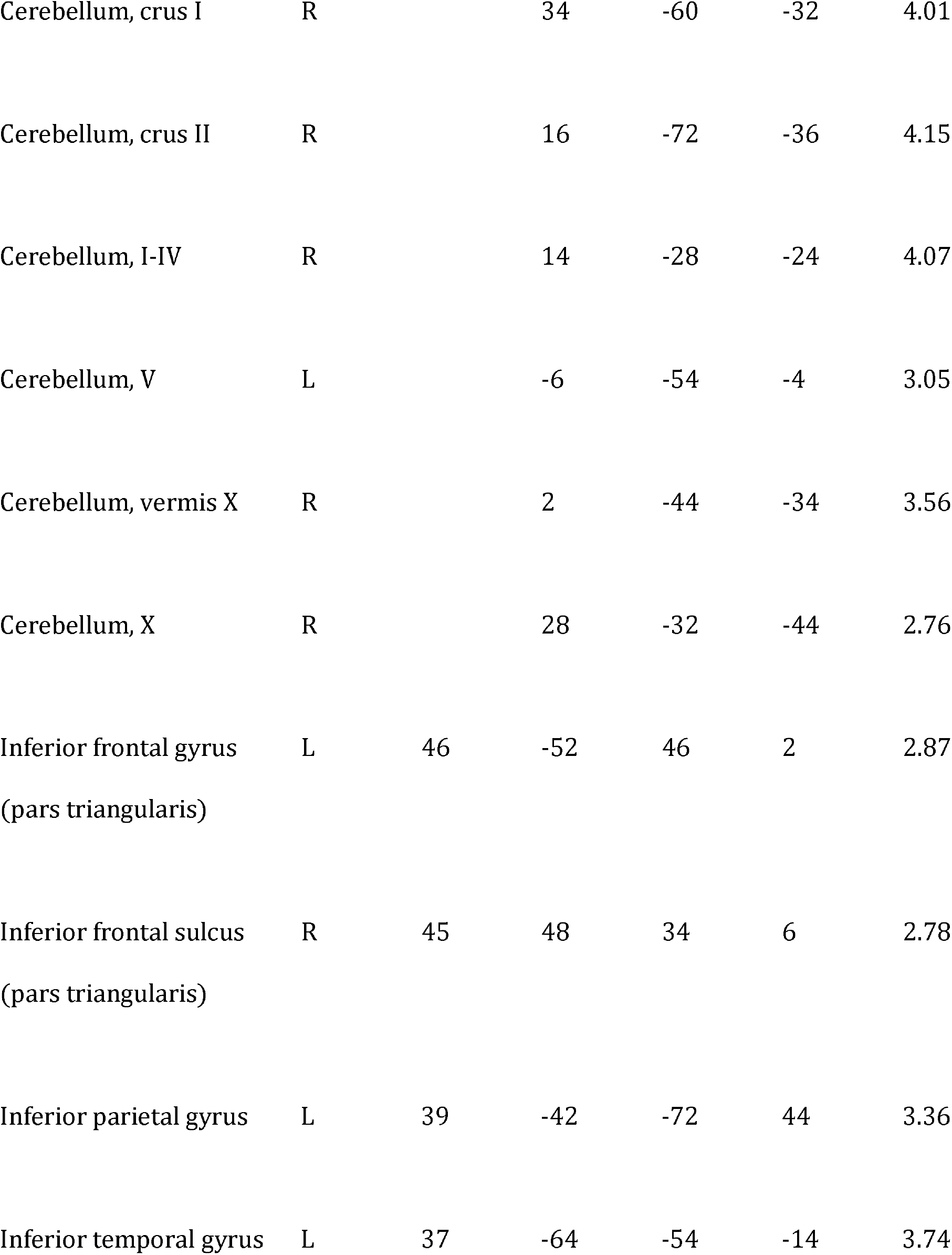
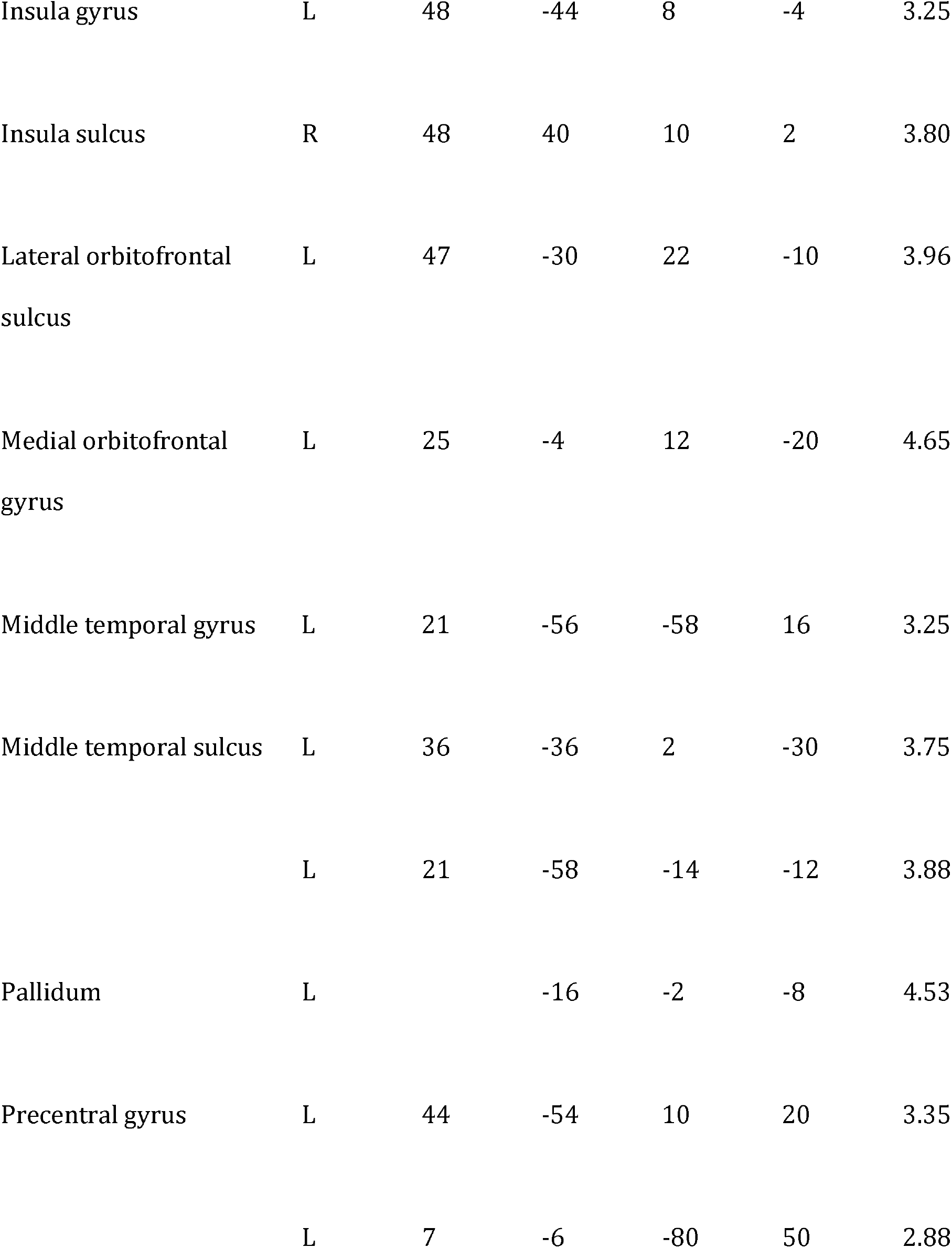
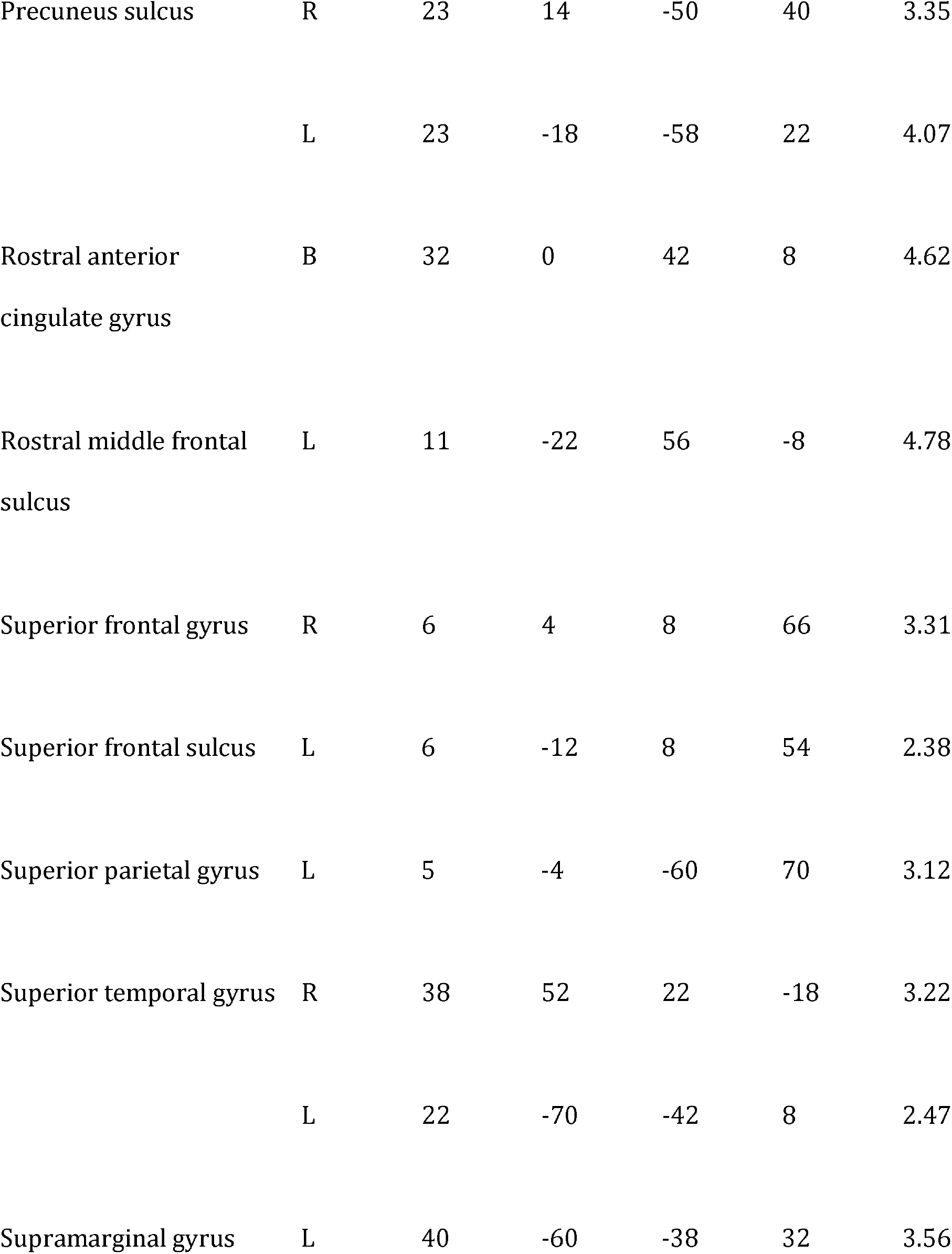
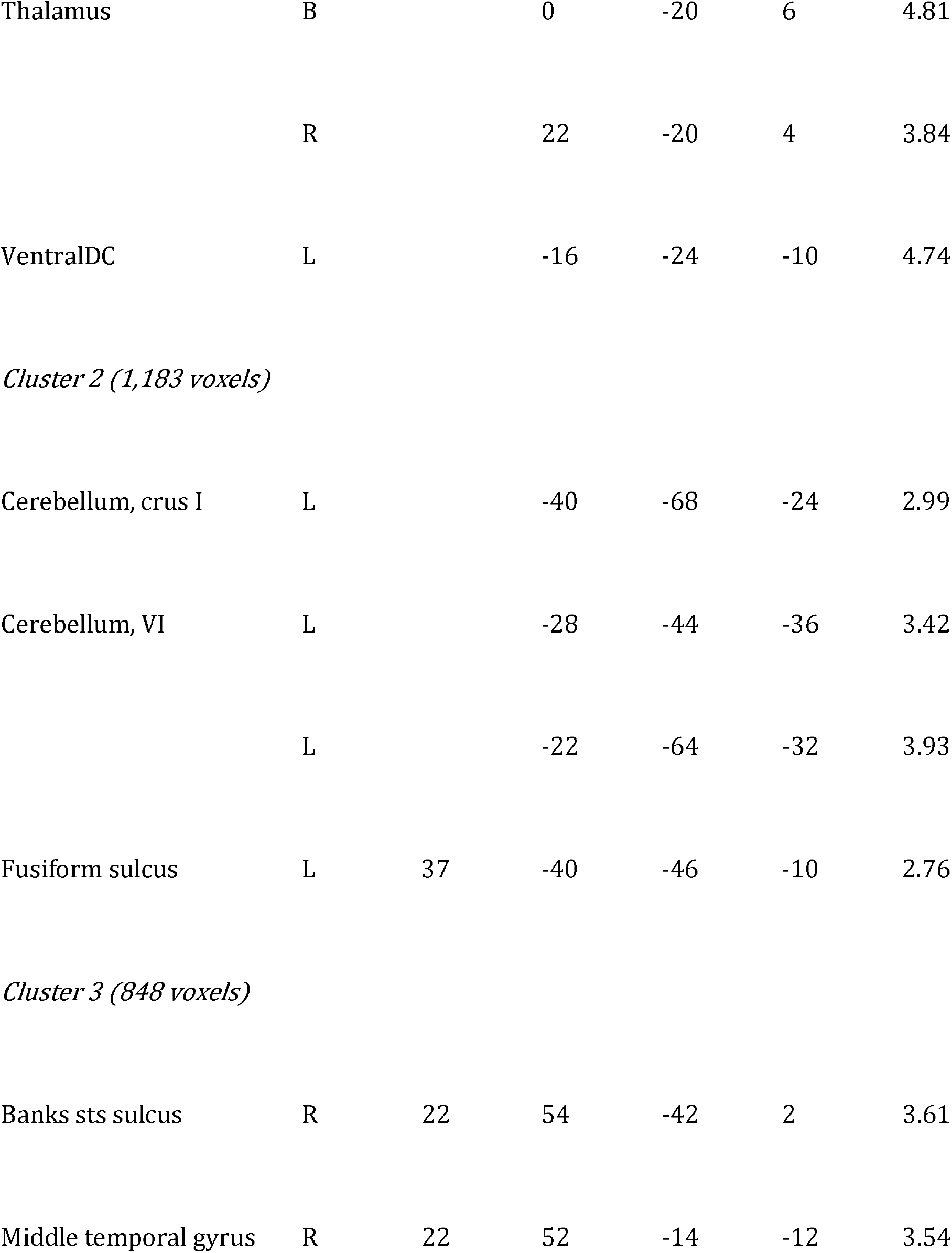
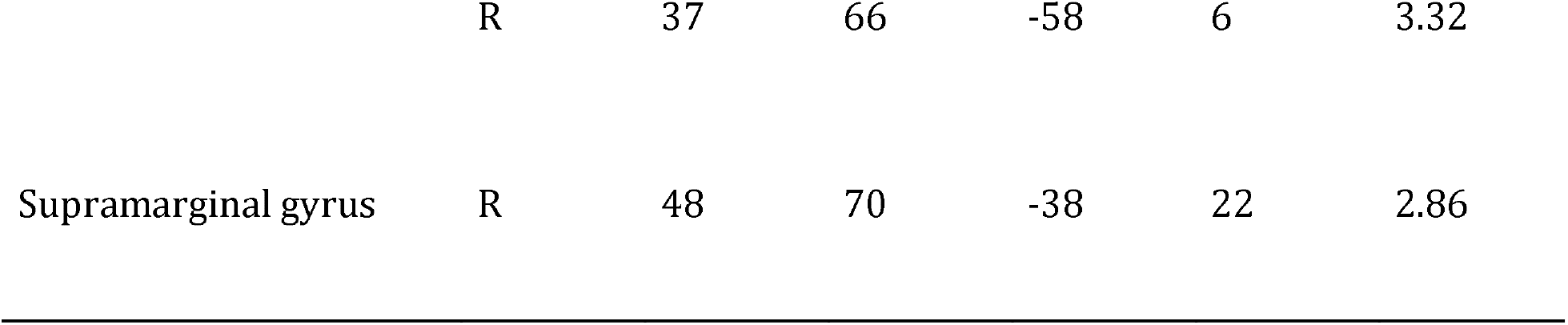
Listed are the peaks of significant brain activity during the Passive Viewing task for faces associated with (bio)graphical versus (phys)ical information. Peaks were found using AFNI's 3dExtrema by setting a voxel-level threshold of Z > 1.96 (cluster corrected, p < 0.05) and a minimum distance of 20 mm between peaks. Abbreviations: H, hemisphere; B, bilateral; R, right; L, left; BA = Brodmann Area.

| Brain region | H | BA | x | y | z | Z-Score |
| --- | --- | --- | --- | --- | --- | --- |
| Caudal anterior cingulate gyrus | R | 24 | 2 | 26 | 24 | 4.29 |
| Cerebellum, crus I | R |  | 34 | -60 | -32 | 4.01 |
| Cerebellum, crus II | R |  | 16 | -72 | -36 | 4.15 |
| Cerebellum, I-IV | R |  | 14 | -28 | -24 | 4.07 |
| Cerebellum, V | L |  | -6 | -54 | -4 | 3.05 |
| Cerebellum, vermis X | R |  | 2 | -44 | -34 | 3.56 |
| Cerebellum, X | R |  | 28 | -32 | -44 | 2.76 |
| Inferior frontal gyrus<br>(pars triangularis) | L | 46 | -52 | 46 | 2 | 2.87 |
| Inferior frontal sulcus<br>(pars triangularis) | R | 45 | 48 | 34 | 6 | 2.78 |
| Inferior parietal gyrus | L | 39 | -42 | -72 | 44 | 3.36 |
| Inferior temporal gyrus | L | 37 | -64 | -54 | -14 | 3.74 |
| Insula gyrus | L | 48 | -44 | 8 | -4 | 3.25 |
| Insula sulcus | R | 48 | 40 | 10 | 2 | 3.80 |
| Lateral orbitofrontal<br>sulcus | L | 47 | -30 | 22 | -10 | 3.96 |
| Medial orbitofrontal<br>gyrus | L | 25 | -4 | 12 | -20 | 4.65 |
| Middle temporal gyrus | L | 21 | -56 | -58 | 16 | 3.25 |
| Middle temporal sulcus | L | 36 | -36 | 2 | -30 | 3.75 |
|  | L | 21 | -58 | -14 | -12 | 3.88 |
| Pallidum | L |  | -16 | -2 | -8 | 4.53 |
| Precentral gyrus | L | 44 | -54 | 10 | 20 | 3.35 |
|  | L | 7 | -6 | -80 | 50 | 2.88 |
| Precuneus sulcus | R | 23 | 14 | -50 | 40 | 3.35 |
|  | L | 23 | -18 | -58 | 22 | 4.07 |
| Rostral anterior cingulate gyrus | B | 32 | 0 | 42 | 8 | 4.62 |
| Rostral middle frontal sulcus | L | 11 | -22 | 56 | -8 | 4.78 |
| Superior frontal gyrus | R | 6 | 4 | 8 | 66 | 3.31 |
| Superior frontal sulcus | L | 6 | -12 | 8 | 54 | 2.38 |
| Superior parietal gyrus | L | 5 | -4 | -60 | 70 | 3.12 |
| Superior temporal gyrus | R | 38 | 52 | 22 | -18 | 3.22 |
|  | L | 22 | -70 | -42 | 8 | 2.47 |
| Supramarginal gyrus | L | 40 | -60 | -38 | 32 | 3.56 |
| Thalamus | B |  | 0 | -20 | 6 | 4.81 |
|  | R |  | 22 | -20 | 4 | 3.84 |
| VentralDC | L |  | -16 | -24 | -10 | 4.74 |
| *Cluster 2 (1,183 voxels)* |  |  |  |  |  |  |
| Cerebellum, crus I | L |  | -40 | -68 | -24 | 2.99 |
| Cerebellum, VI | L |  | -28 | -44 | -36 | 3.42 |
|  | L |  | -22 | -64 | -32 | 3.93 |
| Fusiform sulcus | L | 37 | -40 | -46 | -10 | 2.76 |
| *Cluster 3 (848 voxels)* |  |  |  |  |  |  |
| Banks sts sulcus | R | 22 | 54 | -42 | 2 | 3.61 |
| Middle temporal gyrus | R | 22 | 52 | -14 | -12 | 3.54 |
|  | R | 37 | 66 | -58 | 6 | 3.32 |
| Supramarginal gyrus | R | 48 | 70 | -38 | 22 | 2.86 |

### Classification

#### Questions Task

We examined whether biographical knowledge might also be reflected in spatial patterns of brain activity within the vATL and potentially within other face-selective ROIs for the Questions task. We found that patterns of activity significantly discriminated between conditions in all face-selective ROIs except the R IOG (*ps* < 0.05; Figure 4a). The highest classification accuracies for the Questions task were present in the left mFus (64.4 ± 2.4% [mean ± standard error]), left vATL-p (64.0 ± 1.6%), and left vATL-a (61.0 ± 1.9%). We compared the amount of shared and unique information present in our face-selective ROIs by measuring how much additional unique information each ROI contributed over and above the other ROIs (within each hemisphere). In the Questions task, we found significant unique information for discriminating between the biographical and physical conditions in the left mFus (relative accuracy 2.5 ± 1.2%) and left vATL-p (relative accuracy 2.6 ± 1.5%) (*ps* < 0.05; Figure 4b). Thus, while other ROIs (such as the left vATL-a) had information in their pattern of brain activity differentiating the two conditions, this information was not unique to those ROIs as it was also present in the left mFus and vATL-p.

**Figure 4:**
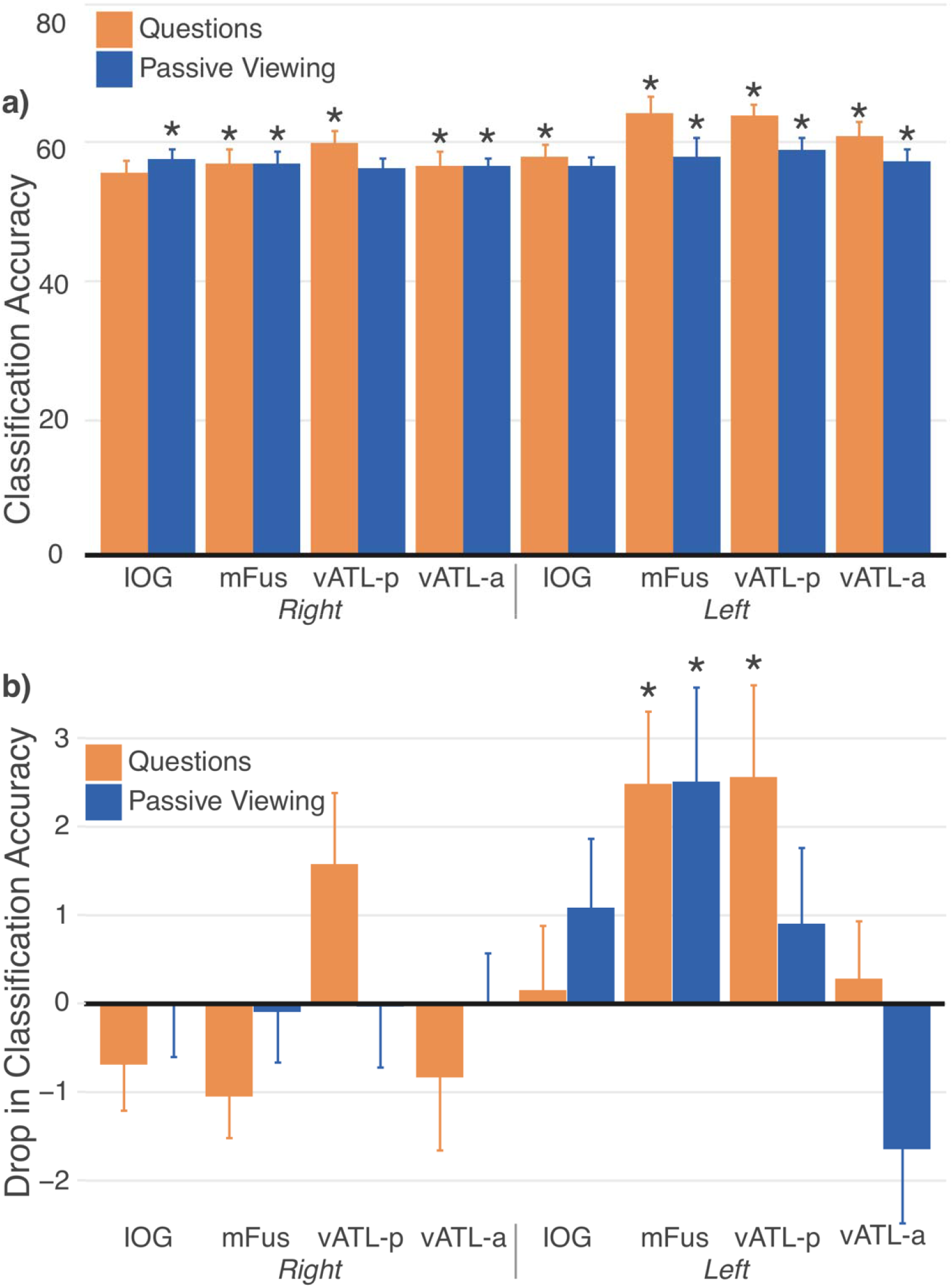
(a) Pattern classification accuracy for discriminating between the biographical and physical conditions (y-axis) is shown for each face-selective ROIs (x-axis). Orange bars indicate the Questions task and dark blue bars indicate the Passive Viewing task. Regions with significant results are indicated with an asterisk. (b) The relative pattern classification accuracy (y-axis) is shown for each ROI (x-axis). For each hemisphere, the classification of all the ROIs (full model) was compared to all the ROIs except one (left-out model; full – left-out). High relative accuracies indicate the left-out ROI contributed additional information to discriminating between the biographical and physical conditions over and above the other ROIs. Negative relative accurac es indicate that the classifier performed better without the ROI in the model (i.e., adding the ROI to the model made the classifier perform worse).

Since the classification accuracy indicates the presence of an effect but not its direction, we do not know if the unique information reflects person knowledge per se (Bio > Phys) or physical face features (Phys > Bio). To better understand the nature of the representations in the left mFus and vATL-p, we examined the voxelwise feature weights for predicting the biographical versus physical condition in a multivariate model that combined all regions within each hemisphere together (Figure 5). We divided voxels into two groups, one with feature weights associated with the biographical condition (i.e., where an increase in activity is associated with a higher likelihood of the biographical condition) and another with feature weights associated with the physical condition (i.e., where an increase in activity is associated with a higher likelihood of the physical condition). We found that the left vATL-p had significantly more feature weights associated with the biographical condition (3.1 ± 0.6) than the physical condition (1.6 ± 0.5; *p* < 0.05)^1^. In contrast, the left mid-fusiform had a comparable number of features associated with the biographical (2.6 ± 0.6) and physical conditions (2.5 ± 0.7; *p* = 0.84). Since the feature weights of sparse multivariate models can be unstable and vary considerably with the addition or removal of voxels (Xu and Caramanis, 2011), we re-estimated the weights for each voxel in a univariate model and found similar results. The feature weights suggest that for the Questions task the information in the left vATL-p is specialized for the biographical condition while the information in the left mFus is equally relevant for both the biographical and physical conditions.

**Figure 5:**
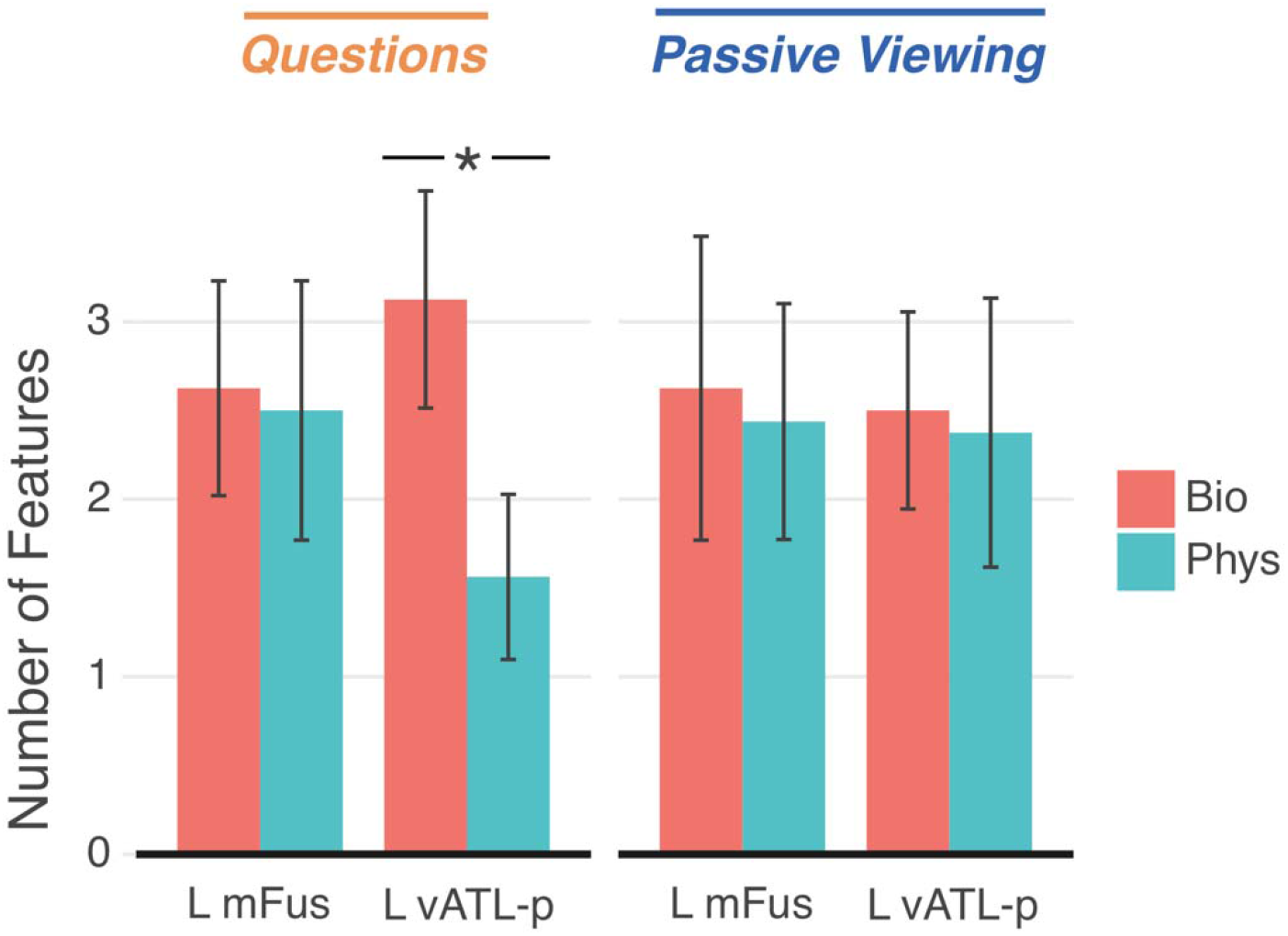
Using a multivariate regression, the number of feature weights (y-axis) associated with the biographical (red) or physical (blue) condition are shown in the left mFus and vATL-p for the Questions and Passive Viewing tasks (x-axis). Feature weights were calculated using a regularized logistic regression with an elastic-net penalty that included voxels within the same hemisphere. Error bars reflect standard error of the mean, and an asterisk indicates a significant difference (p < 0.05) between the two conditions.

#### Passive-Viewing Task

We also examined spatial patterns of activity for the Passive Viewing task. We found that patterns of activity significantly discriminated between conditions in all face-selective ROIs except the R vATL-p and L IOG (ps < 0.05; Figure 4a). The highest classification accuracies for the Passive-Viewing task were present in the left mFus (58.1 ± 2.8% [mean ± standard error]), left vATL-p (59.2 ± 2.8%), and left vATL-a (57.4 ± 1.8%). When separating the unique from shared information, we found significant unique information for discriminating between the biographical and physical conditions only in the left mFus (relative accuracy 2.5 ± 1.6%) (*ps* < 0.05; Figure 4b). Next, we examined the feature weights to better understand the nature of the information in the left mFus (i.e., does it reflect activity related to the biographical or physical condition; Figure 5). The left mFus had a comparable number of features associated with the biographical condition (2.6 ± 0.9) and physical condition (2.4 ± 0.7; *p* = 0.75). For consistency with the Questions task, we also examined the feature weights in the left vATL-p for the Passive-Viewing task. As with the left mFus, the left vATL-p had a comparable number of features associated with the biographical condition (2.5 ± 0.6) and physical condition (2.4 ± 0.8; *p* = 0.79). The feature weights suggest that for the Passive-Viewing task the information in both the left mFus and vATL-p is equally relevant for the biographical and physical condition.

## Discussion

This study investigated whether person knowledge for faces is a localized process restricted to the vATL or a distributed process spread across face-selective regions. Prior work on person knowledge has been inconclusive, providing support for either a localized, modular (Nobre et al. 1994; Rotshtein et al. 2005; Von Der Heide et al. 2013; Anzellotti et al. 2014) or distributed (Haxby et al. 2001; Verosky et al. 2013; Axelrod and Yovel 2015; Yang et al. 2016) neural architecture. Our findings suggest a mixture of the two models can explain the difference in neural signals for faces associated with biographical versus physical information. We first found distributed representations across the bilateral ventral surfaces of the occipital and temporal lobes when examining each region (excepting the rIOG) in isolation, which was true for both the Questions (explicit) and Passive Viewing (implicit) tasks. However, when we combined the regions into one classification model, we found much of this biographical/physical information was shared or redundantly represented among regions. After removing the shared information between regions, we found the unique information to be localized in the left hemisphere. Unique information was found in the left mid-fusiform for both tasks but reflected a mix of biographical and physical information about a face. In contrast, unique information was found in the left vATL-p only when person knowledge was explicitly retrieved (Questions task), and in this context the representations in the left vATL-p were specialized for biographical and not physical information. Thus, our novel findings reconcile a debate in the localization of person knowledge for faces and suggest that the left mid-fusiform represents aspects of face processing not specific to person knowledge (e.g., perception) that can be implicitly retrieved while the left vATL-p specifically represents person knowledge associated with a face that must be explicitly retrieved.

### vATL a module for biographical information about faces

Our findings support the role of the vATL in “linking perceptual representations of individual identity with person-specific semantic knowledge” (Collins and Olson 2014). In the present study, faces associated with biographical compared to physical facts produced more unique multivariate information and greater univariate activity in the vATL-p but not in other face-selective regions (Figure 3 and Supplementary Figures 1-2 [https://doi.org/10.5281/zenodo.3700452]). Unlike more typical studies that examine familiar faces, our task design controlled for the amount of exposure to each identity between the Bio and Phys conditions. Consequently, our results add to the role of the vATL in face processing, suggesting that it responds to person-specific knowledge regardless of the perceptual familiarity with the face.

We identified two regions within vATL: the vATL-a and the vATL-p. Only the left vATL-p was significantly associated with person knowledge. The vATL-a is typically activated for familiar versus novel faces (Von Der Heide, 2013) and for faces associated with conceptual knowledge (Barense et al., 2011; Eifuku et al., 2011; Ross and Olson, 2011; Nieuwenhuis 2012). Several possibilities exist for finding unique representations of person knowledge in the left vATL-p and not the more anterior vATL-a. Signal drop-out could have been a factor. The vATL-a is known to suffer from severe fMRI signal drop-out due to magnetic susceptibility caused by the ear canal (Carr et al., 2010; De Panfilis and Schwarzbauer, 2005; Devlin et al., 2000; Gorno-Tempini et al., 2002; Kriegeskorte et al., 2007; Leopold et al., 2006; Olman et al., 2009; Rajimehr et al., 2009). To compensate, prior work that has found robust vATL activity has used pulse sequences optimized for ATL coverage and sensitivity, which was not done here (Von Der Heide et 2013, Axelrod et al, 2013). This could result in a larger signal drop-out for the vATL-a than vATL-p ROI and explain the null results in the vATL-a. Although we found lower SNR in the left vATL-a than the left vATL-p corroborating this possibility, the difference was not significant (Supplementary Figure 1 [https://doi.org/10.5281/zenodo.3700452]).

Another factor could be differences in our task design. Since the task required participants to memorize only three facts associated with a face, it could be that retrieval of a specific and limited number of person attributes involves the left vATL-p. In contrast, more anterior portions of the vATL might be relevant for binding rich sets of facts for an individual such as a famous face.

The final and perhaps most important factor is that prior work has not compared representations between regions. When we examined ROIs in isolation, we found significant classification accuracy for the left vATL-a, a result consistent with prior work. However, when comparing the representations of the left vATL-a with other regions, we found the information in the left vATL-a was not unique. Our results then suggest that the more anterior portions of the vATL may have information for person knowledge (consistent with prior work) but this information is not unique and instead shared with the vATL-p.

### Lateralization of biographical information about faces

We found a left-lateralization of biographical information for faces for univariate measures of brain activity and multivariate classification accuracy (Figure 3, Supplementary Figures 1-2 [https://doi.org/10.5281/zenodo.3700452]). This differentiates our findings from the right lateralized brain responses found when examining non-verbal aspects of face processing (Gazzaniga and Smylie 1983; De Renzi et al. 1994; Le Grand et al. 2003) or representations of face identity in the right vATL (Kriegeskorte et al. 2007; Nestor et al. 2011; Verosky et al. 2013). In contrast, our task had a greater verbal component since we compared biographical facts (a form of semantic knowledge) versus physical facts (a form of visual knowledge) associated with individual faces that were familiarized through training. Similarly, left-lateralized activations in the vATL have been found for personally familiar and famous individuals, while novel faces have activated the right vATL (Von Der Heide et al. 2013). Furthermore, human lesion work suggests that the left ATL is related to processing semantic information associated with individuals as in our study (e.g., proper names), whereas the right ATL is associated with processing visual information related to faces (Gainotti 2007; Brambati et al. 2010). We note that when examining the average level of activity across our two conditions (i.e., summing across biographical and physical), we found the temporal response in the mid-fusiform had visibly greater activity in the right hemisphere than left hemisphere (Figure 4), consistent with the claim that overall face-processing is right lateralized in the mid-fusiform (Kanwisher et al. 1997; McCarthy et al. 1997).

### Left mid-fusiform sensitive to non-visual information

Our findings support a role of the mid-fusiform (our mFus ROI) in person knowledge. Typically, the mid-fusiform has been associated with unimodal visual information relevant to faces but several studies suggest it may respond to other sensory modalities including semantic information relevant to person knowledge (von Kriegstein, Kleinschmidt, Sterzer, & Giraud, 2005; Mahon, Anzellotti, Schwarzbach, Zampini, & Caramazza, 2009; Van den Hurk, Gentile, & Jansma, 2011). For example, patterns of activity for faces in the mid-fusiform were grouped based on the biographical information learned for each identity (Verosky et al., 2013). Even in the absence of face stimuli, the mid-fusiform responds differently to words describing categories of objects versus people, showing sensitivity to semantic context (van den Hurk et al., 2011).

We also examined semantic content in the biographical condition, albeit associated with faces. When examining the pattern information in the mid-fusiform, we found an even distribution of voxels associated with biographical and physical information of a face (Figure 4). For the physical condition, we expected greater perceptual processing since participants explicitly attended to and learned about physical face features. Consequently, if information was predominantly perceptual, we would have expected more voxels responding with higher activity to the physical condition. These results indicate that both perceptual information (physical condition) and semantic information (biographical condition) might be present in the mid-fusiform. However, unlike the left vATL-p, the feature weights in the left mid fusiform were not affected by explicitly asking participants to retrieve the biographical facts. This suggests that the left mid-fusiform may not be involved in the explicit and conscious retrieval of person knowledge but in an earlier and automatic process relevant to person knowledge.

### Do classification results support distributed processing of biographical knowledge?

In face-selective ROIs, multivariate pattern analyses revealed a distributed representation for biographical knowledge of faces, whereas univariate analyses indicated a localized representation wherein biographical information resides only in the vATL. One explanation for such differences in the two methods is increased sensitivity for multivariate analyses compared to univariate (Jimura and Poldrack 2012; Davis and Poldrack 2014). Multivariate approaches have been shown to provide more power to detect an effect and to be more robust to noise than univariate analyses (Vickery et al. 2011; Jimura and Poldrack 2012). This increase in power may reflect the fact that, unlike univariate analyses, multivariate analyses are sensitive to voxel-level variability within subjects and ignore subject-level variability in mean activations (Davis and Poldrack 2014). Multivariate analyses tend to find optimal weightings of each voxel when predicting task activity for each subject. Consequently, even when there is no regionally coherent and/or robust response in univariate responses of certain brain areas (e.g., face-selective regions other than the vATL), there can still be information found in multivariate patterns of activity.

Demonstrating high classification accuracy across face-selective regions, however, does not on its own reflect distributed processing of biographical information for faces, and further examination of the information content is required (Davis and Poldrack 2014). One issue is that multivariate classification analyses can use any information to discriminate the biographical and physical conditions, which may include differences between conditions irrelevant to biographical knowledge such as task difficulty or reaction-time. Even when only task-relevant information is used by a classifier, different brain regions may have high classification accuracy but contain the same information content as another region (i.e., information is shared between regions), which would invalidate any claims of a distributed neural architecture (i.e., each region contributes unique information to a task). Removing the shared information across regions allows the identification of those regions that contribute additional information over and above other regions for a particular task.

To measure the potential effects of shared, or redundant, information, we used a procedure that we previously developed to remove shared information content between regions (Shehzad and McCarthy, 2018). We measured classification accuracy in each region while removing the covariation with other regions. Any remaining information would be unique to that region. If biographical versus physical information is distributed, then we would expect that even after removing shared information between regions, there would be residual unique information in each region. This was not found. Rather, our results support a more localized model in which unique information for biographical knowledge was localized to the left mFus and left vATL-p. Thus, there was a large degree of shared information between regions, which resulted in the appearance of distributed representations when examining classification accuracy in each region individually.

Given the major portion of shared information among regions, what is the nature of the shared information identified here, and its role in neural function? We have previously discussed several possibilities (Shehzad et al., 2018). Briefly, shared information might be redundant and reflect an epiphenomenon of network connectivity (for example, much like sound information can spread through a space). In contrast, the shared information might be functional and could reflect inputs from other regions that are then used for local computations. Finally, shared information could reflect preparation or predictive coding (for example, biographical information about a person could prime visual activity of that person's face).

### Connections to other methods?

Our approach has similarities to Bayesian network analyses and graphical causal models (Mumford and Ramsey, 2014). These approaches can also estimate the unique versus shared information within a network by resolving the direction and propagation of information flow. However, the nature of the information is often unclear with these connectivity approaches. Our approach estimates information relative to an outcome measure (e.g., biographical vs physical faces), specifically we test the relationship between a region and outcome conditional on other regions. In this way, we can interrogate different types of information (e.g., low-level vs high-level visual information) by examining different outcome measures with our classifier. Here, we focused on comparing representations for faces associated with biographical versus physical information but other work could examine additional aspects of person knowledge such as the content of the biographical (e.g., name) information.

In this study, the use of an independent probabilistic atlas for the ROIs allowed for us to generalize face-selective regions across participants. However, these group-based ROIs may overlook subject-specific activity for face processing as in face localizers and may weaken our results. Future work may consider combining the use of a probabilistic atlas with face localizers to improve reproducibility. In work using resting-state functional connectivity, it has been shown that the use of a group-average functional connectivity map as a prior can improve the reproducibility of individual connectivity maps (Shou et al., 2014). A similar approach could be used to improve the reliability of an individual face localizer by using the group average as a prior.

Our work has a number of limitations. By design, our classification model only examined regions along the ventral surface known to be face-specific. This prevents us from making broader claims about the role of regions beyond those in our model. Since our explicit goal was to adjudicate the roles of the mFus and vATL in person knowledge for faces, understanding the role of other regions was beyond the scope of the present paper. In addition, there were attentional differences between the biographical and physical conditions. Due to the greater difficulty of recalling biographical facts (physical facts were apparent on the face), differences in activity between the two conditions could reflect greater attentional demands in the biographical condition. For example, a subsequent study might include a third condition that requires recalling facts unrelated to a face to control for attentional demands.

## Conclusion

We asked participants to view faces previously associated with biographical facts compared to equally perceptually familiar faces associated with observable physical facts. Robust univariate activations were observed in the vATL for biographical knowledge, suggesting a modular response. Significant classification accuracy between the biographical and physical conditions was found across face-selective regions, suggesting a distributed response. However, when we removed shared information present between regions, we observed a more localized response in the left mFus and vATL-p. Furthermore, unique information for biographical versus physical facts was only found for in the left vATL-p (and not in the left mFus) when biographical facts were explicitly retrieved. These results suggest that specialized representations for person knowledge exist in the left vATL-p and require explicit retrieval, while representations for face processing not specific to person knowledge exist in the left mFus and can be automatically retrieved. Taken together, our findings support a model of face processing where the mid-fusiform is associated with an initial perceptual analysis of a face composed of a fine-grained analysis of physical details, whereas the ventral anterior temporal lobe contains neural populations that encode biographical information.

## Acknowledgements

This work was supported by the National Institute of Mental Health (MH-005286 to G.M.). We would like to thank Eunjoo with assistance in reading and editing the comments/manuscript, as well as Magdaleno Mora and Audrey Luo for their assistance with data collection.

## Footnotes

1 The regularization procedure removes voxels with redundant information. Hence, the total number of features can be taken as a measure of the total amount of information for biographical or physical information present in a ROI (Diedrichsen et al., 2013). Since the values (non-zero features) are much lower than the total number of voxels in the ROI, this suggests that there are a small number of undeg features associated with the task in each region.

