## Supplementary Materials for "Localized and Distributed Representations of Person Knowledge for Faces"

### Supplementary Methods

#### Regions of Interest – Control Analyses

Using the Atlas of Social Agent Perception (Engell and McCarthy 2013), we identified two activation peaks based on the average of two probability maps (house > face and scene > face). The peaks were selected to be in regions not expected to be face-selective and included the left lingual gyrus (Ling; X = -28, Y = -46, Z = -8; BA 37) and the pre-central gyrus (PrC; X = 2, Y = -22, Z = 56; BA 6). We used the same procedure to generate the probabilistic maps, activation peaks, and ROIs for these two control ROIs as in the main paper for the face-selective ROIs.

### Supplementary Results

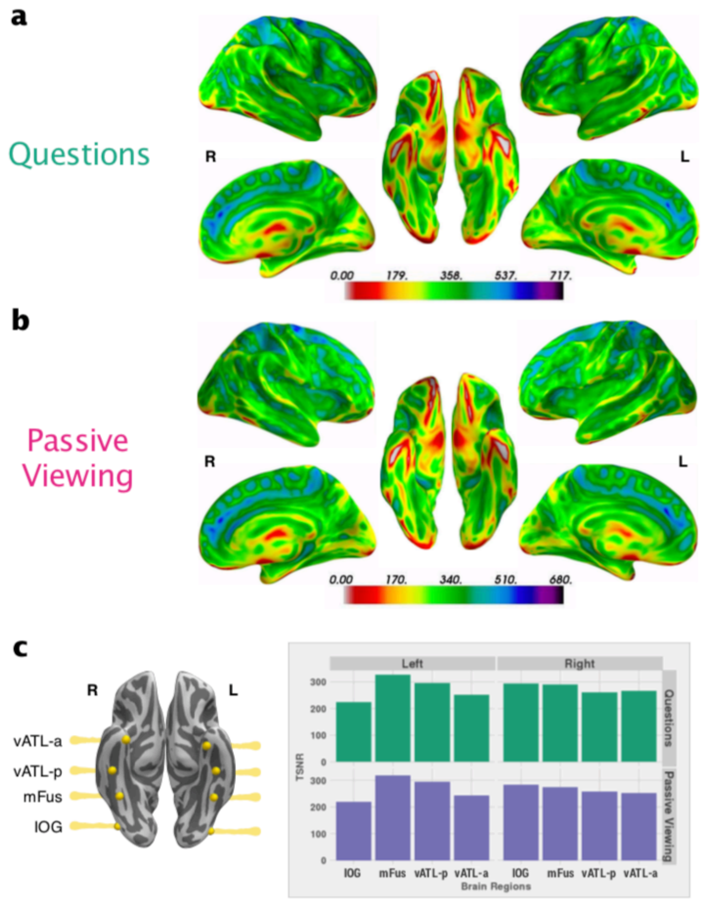


**Supplementary Figure 1.** Temporal signal-to-noise ratio (TSNR) maps showing image quality of the functional data. At each voxel, TSNR was calculated by dividing the mean signal intensity of the preprocessed functional time-series by the standard deviation of the residual time-course from the task analysis. TSNR values are overlaid on the fsaverage surface using a spectral color gradient for the (a) Questions task and (b) Passive Viewing task. Regions shown in the ventral surface view without any value (no color) reflect voxels that have no signal coverage. (c) Face-selective ROIs shown on the ventral surface were used to calculate the average TSNR values (y-axis) in each of the face-selective ROIs (x-axis) for each task and are displayed as a bar plot. Simulations have shown that a TSNR of at least 40 is needed to reliably detect effects between conditions in fMRI data (Murphy et al. 2007). Note that all the face-selective ROIs including the vATL-a far exceed this threshold with a minimum TSNR of

~200. However, on the ventral surface, signal drop out shown as red can be observed proximal to the ear canal.

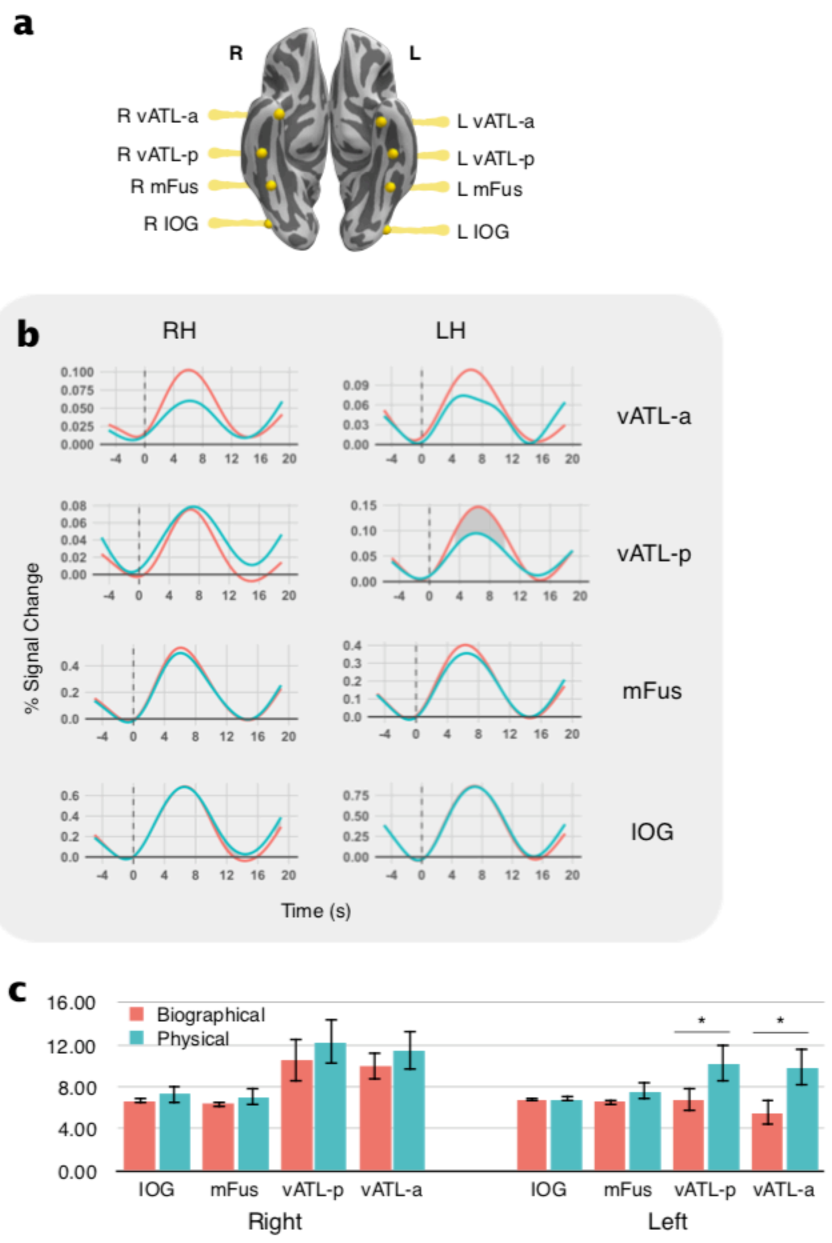


**Supplementary Figure 2:** (a) Face-selective ROIs rendered on the fsaverage surface. (b) Average evoked response using smoothed splines when viewing faces in the biographical (red) or physical (blue) condition for the Questions task. The x-axis indicates time in seconds (dashed line = onset of the face), and the y-axis indicates the percent signal change relative to baseline (mean signal 0-2s before onset). (c) Average peak latency in seconds across subjects (y-axis) is given for each of the ROIs (x-axis). ROIs with significant differences between condition are represented with a * ($p < 0.05$).

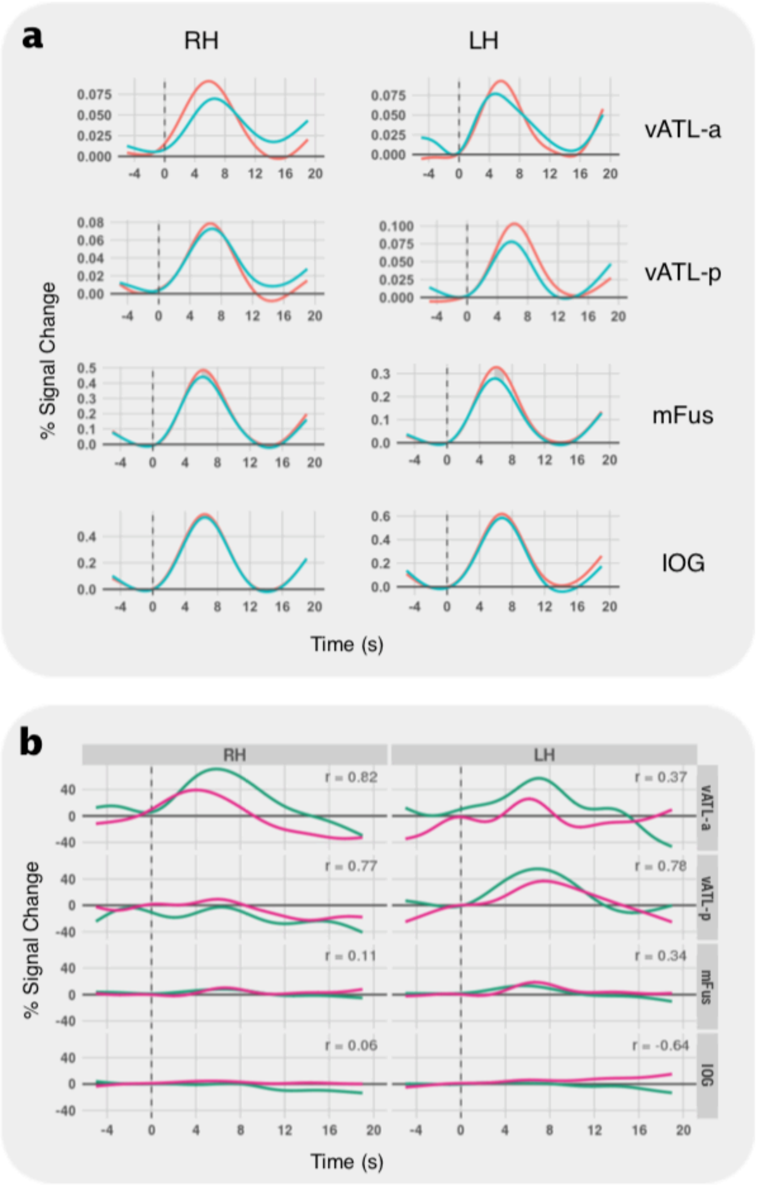


**Supplementary Figure 3:** (a) Average evoked response using smoothed splines when viewing faces in the biographical (red) or physical (blue) condition for the Passive Viewing task. The x-axis indicates time in seconds (dashed line = onset of the face), and the y-axis indicates the percent signal change relative to baseline (mean signal 0-2s before onset). (b) Temporal difference curves between the Bio and Phys average evoked responses for the Question (green)

and Passive Viewing (purple) tasks. The Spearman rho between the temporal difference curves for the two tasks for each region is shown.

#### Classification – Control Analyses

We ran additional control analyses in the left hemisphere to test if our if non-face-selective regions contained unique or shared information for person knowledge associated with faces. We examined information in our left hemisphere ROIs (IOG, mFus, vATL-p, vATL-p) as well as in non-face-selective regions (Ling, PrC) for both the Questions and Passive Viewing tasks. When measuring the classification accuracy for each region separately, the results for the face-selective regions will be the same as our prior analysis (i.e., in Fig 5). However, when putting all six regions together into one model, the classification results for the face-selective regions will change depending on the amount of shared information removed by the addition of the other regions.

*Questions Task.* We found that patterns of activity significantly discriminated between conditions in our two non-face-selective regions (*ps* $< 0.05$; Supp Fig 4a). The classification accuracies in the control regions were 58.1 ± 5.6% (mean ± standard error) for the left Ling and 58.2 ± 7.4% for the PrG. We found a different pattern of results when we measured the unique information that each control ROI contributed over and above the other ROIs. There was no significant unique information for discriminating between the biographical and physical conditions in the left Ling (relative accuracy 0.19 ± 0.57%, $p = 0.42$) and PrG (relative accuracy 0.26 ± 0.43%, $p = 0.40$; Supp Fig 4b). When examining the pattern of unique information for our face-selective ROIs, the results remained the same for the left hemisphere whether or not the control ROIs were included in the model. There was significant unique information in the left mFus (relative accuracy 2.51 ± 1.37%) and left vATL-p (relative accuracy 2.55 ± 1.86%) (ps $< 0.05$, Supp Fig 4b) while the information in other face-selective ROIs (left IOG and left vATL-p) was not unique. These findings confirm that person knowledge for faces in the Questions task was localized in the left mFus and vATL-p and distributed across other regions including those there were non-face-selective.

*Passive-Viewing Task.* We also examined spatial patterns of activity for the Passive Viewing task using our control ROIs. As in the Questions task, we found that patterns of activity significantly discriminated between conditions in all ROIs including non-face-selective regions ($ps < 0.05$; Supp Fig 4a). The classification accuracies in the control regions were 57.8 ± 6.9% for the left Ling and 59.9 ± 6.9% for the PrG. However, when we separated the unique from shared information, the two control regions did not have significant unique information for discriminating between the biographical and physical conditions ($ps > 0.6$; Supp Fig 3b). In the face-selective regions, we found marginally significant unique information only in the left mFus (relative accuracy 2.1 ± 1.6%, $p = 0.06$), which is a similar result to our analysis with only the face-selective regions.

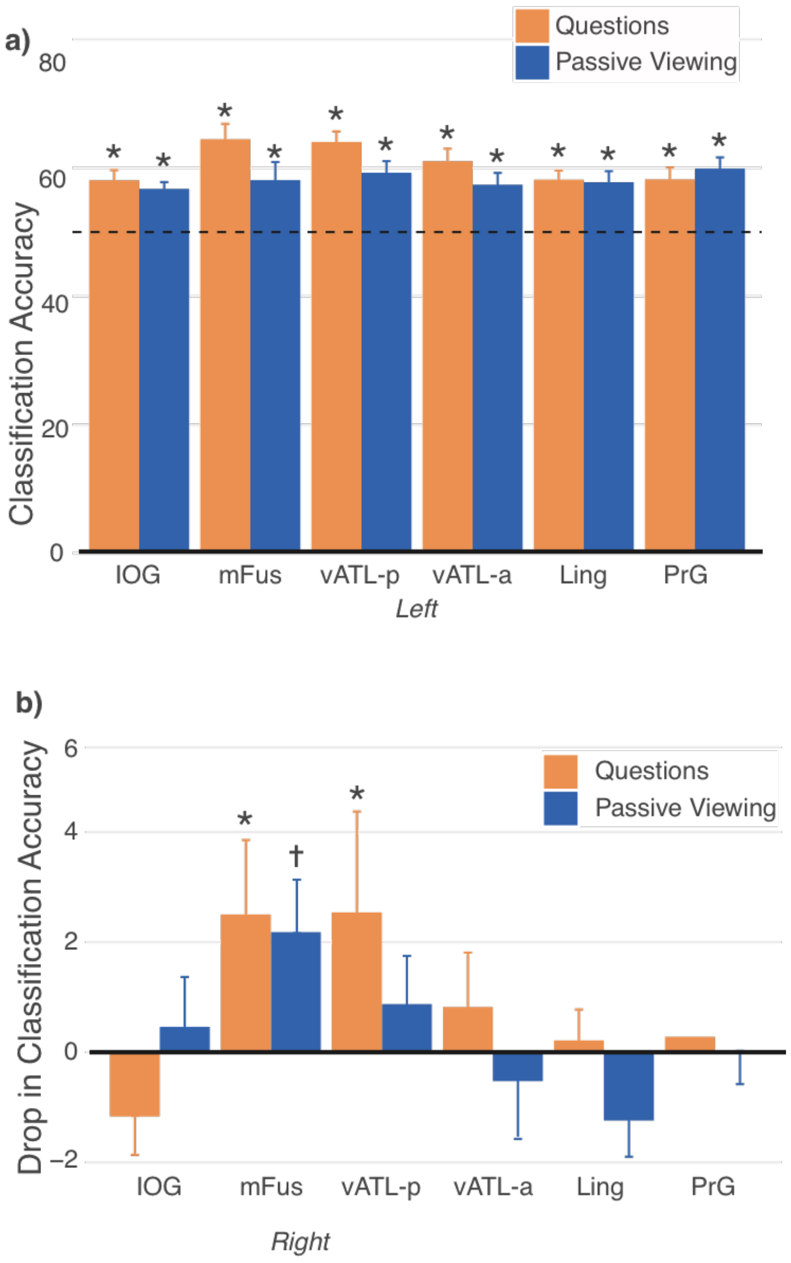


**Supplementary Figure 4:** (a) Pattern classification accuracy for discriminating between the biographical and physical conditions (y-axis) is shown for each face-selective ROIs in the left hemisphere and two control ROIs (x-axis). Orange bars indicate the Questions task and dark blue bars indicate the Passive Viewing task. Error bars reflect standard error of the mean. Regions with significant results are indicated with an asterisk ($p < 0.05$). Significance was determined with a permutation test (using 500 permutations). (b) The relative pattern classification accuracy (y-axis) is shown for each ROI (x-axis). For each hemisphere, the classification of all the ROIs (full model) was compared to all the ROIs except one (left-out model; full – left-out). High relative accuracies indicate the left-out ROI contributed additional information to discriminating between the biographical and physical conditions over and above the other ROIs. Negative relative accuracies indicate that the classifier performed better without the ROI in the model (i.e., adding the ROI to the model made the classifier perform worse).
